# Applying Hierarchical Generalized Additive Models to Non-Compartmental Analysis of Pharmacokinetic Data

**DOI:** 10.1101/2023.07.13.548803

**Authors:** Andrew P. Woodward

**Author notes:** Disclosure: The author reports no conflicts of interest. Data and code availability: Data utilized in this manuscript were obtained from published literature and are linked within the text. All R code utilized in this work are available at github.com/APWoodward/NCA_HGAM.

## Abstract

Non-compartmental analysis (NCA) is a popular strategy for obtaining estimates of pharmacokinetic parameters, while requiring both minimal structural assumptions, and limited input by the analyst. As typically applied, its scope and depth are constrained by its statistical simplicity. Embedding the NCA within a hierarchical generalized additive model (HGAM) may facilitate the simultaneous analysis of data from multiple subjects, estimation of covariate effects in one stage, and implementation of censored responses, similarly to the capabilities of nonlinear multilevel models as widely applied in pharmacometrics. HGAM is an interesting extension to multilevel linear models that allows the effects of predictors to be implemented as smooth functions, which has been widely implemented in various disciplines to nonlinear trends, including for longitudinal data.

This approach extends the capability of previous implementations of spline-based methods applied to NCA, within an accessible workflow in open software. Application of HGAM to two example datasets, one describing oral drug administration, and one describing IV and oral drug administration with categorical covariates and censoring, illustrates the overall approach, including parameter estimation, visualization and model checking, and uncertainty quantification. A Bayesian approach to estimation facilitates interpretable expressions of the uncertainty in individual parameters, population parameters, and functions of parameters such as contrasts.

## Introduction

Non-compartmental analysis (NCA) is a popular strategy for statistical analysis of pharmacokinetic (PK) data. Under specific assumptions, the utility of NCA-based estimation of pharmacokinetic parameters is well-established (Gabrielsson and Weiner, 2012). Though population pharmacokinetic methods, comprising parametric nonlinear multilevel models, have become standard methodology in many applications, important roles for NCA remain. NCA offers simple, direct estimation of key pharmacokinetic parameters in cases where limitations of data, knowledge, expertise, or resources constrain the use of more explicit models. Further, NCA is a primary technique in experimental pharmacokinetics, especially in bioequivalence studies, in cases where selection of a structural model is not desired.

The statistical simplicity of NCA, at least within a typical workflow, carries some relevant practical disadvantages. Numeric methods such as trapezoidal rules for the determination of AUC, the key primary parameter estimated in NCA, are reliant on dense sampling, as they have no mechanism to share information between subjects. The sharing of information between subjects (partial pooling), is a key characteristic of population pharmacokinetics (Gelman, Bois and Jiang, 1996), and more generally, multilevel modelling (Greenland, 2000). The numeric methods produce only a point estimate, and do not indicate their uncertainty in the estimated AUC; regardless of the accuracy of the method, the resulting AUC are drawn from finite data and so are not absolutely known. This uncertainty is important for reporting, to reflect the inherent limitation of results of any individual study, but also to the downstream usage of the parameter estimates, as typical in the two-stage approach, for evaluation of bioequivalence for example. Censored data are also not accommodated in the typical workflow for NCA. Recent work highlights the use of various error and imputation models for estimation of population NCA parameters in the presence of left-censoring (Barnett *et al*., 2021), but not individual NCA estimation. In veterinary applications for example, arbitrary decisions such as dropping censored observations are generally abundant (Woodward and Whittem, 2019). In population pharmacokinetics censored observations may be rigorously handled via an appropriate likelihood function (Beal, 2001), but this of course is not present in NCA, which may lead to poor performance (Hughes, Upton and Foster, 2017).

The primary parameter estimates in NCA are integrals; the area under the concentration-time curve (AUC), and the area under the moment concentration-time curve (AUMC), from which other parameters are secondarily derived (Veng-Pedersen, 2001). These are classically obtained using trapezoidal methods, which provide point estimation. Though accurate under suitable conditions (Chiou, 1978; Dunne and King, 1989), they determine only one ‘best’ value, and do not indicate uncertainty. To obtain uncertainty estimation of such integrals, encapsulation of the area estimation within a statistical model, with a suitable likelihood function, is a natural approach. A clear barrier is the specification of the form of the statistical model, given the general principle of non-compartmental modelling to not require a model to be selected by the user. A sensible candidate is a spline-based model, in which the shape of some relationship is a simple smooth function (Tian, Yu and Kim, 2020).

Spline methods can be applied directly to single-level concentration-time data, and examples including cubic splines have been investigated for NCA, with middling results (Purves, 1992). They may also be utilized as elements of a larger statistical model, especially the generalized additive model (GAM; Hastie and Tibshirani, 1986). GAM can be considered a semi-parametric extension of generalized linear models (GLM), in which the relationship between the linear predictor and one or more predictors is expressed by some form of spline (Wood, 2020). In contrast to GLM, which require the investigator to completely specify the form of the relationship between the response variable and all predictors, HGAM allows the form of the effects of one or more of the predictors to be determined from the data during model fitting, allowing for great flexibility. These methods have extensive application in relevant disciplines, such as ecology (Simpson, 2018) and epidemiology (Dominici, 2002), with scope for biomedical applications (Mundo, Tipton and Muldoon, 2022). These methods have a close relationship with other non-parametric techniques such as Gaussian process regression and similar models (Wood, 2020). Current implementations make these models quite accessible. Package ‘mgcv’ (Wood, 2023) implements a variety of computationally efficient HGAM methods (Wood, 2011) using the penalized likelihood, which are also supported in an explicitly Bayesian form in package ‘brms’ (Bürkner, 2017), an interface to the Bayesian programming environment Stan (Carpenter *et al*., 2017).

Various semi-parametric techniques have been widely used in pharmacometrics, generally as components of broader models. Various examples utilize splines to capture the form of covariates (Lai, Shih and Wong, 2006), or to describe time-varying parameters in cases where constant models are insufficient (Li *et al*., 2002). More recently, Gaussian process models were applied to evaluate potentially non-linear maturation effects based on age (Siivola, Weber and Vehtari, 2021). Semi-parametric models for primary analysis of concentration-time functions in pharmacokinetics have been presented sporadically, but appear to have received limited attention. Notably, Park et al., (1997) reported semiparametric modelling using splines, as a convenient and flexible alternative to nonlinear multilevel models, including random effects for between-subject variation. Jullion et al., (2009) described application of Bayesian P-splines, including extensive mathematical details and consideration of priors, with a focus on sparse problems. Most recently, Willemsen et al., (2017) described the application of a specific variant of multilevel spline model to bioequivalence determination with NCA, and nominated several potential advantages, including handling of sparsity and implementation of censoring, which are directly relevant to the HGAM approach evaluated here. Though these evaluations demonstrate substantial utility of a semi-parametric approach to NCA, they are technically sophisticated and demand a relatively high degree of skill to implement.

Broadly, HGAM may facilitate probabilistic, rather than deterministic, estimation in NCA, allowing direct quantification of uncertainty, while aiding the flexibility of analysis. A key goal must be to retain the ease-of-use and data-driven spirit of NCA, relative to other pharmacometric methods which are highly dependent on investigator expertise and nomination of appropriate models. Use of readily available and accessible implementations is thus a key priority. The objective of this manuscript is to explore the application of the hierarchical generalized additive model (HGAM) to NCA, building upon previous applications of semiparametric models for NCA (Park *et al*., 1997; Jullion *et al*., 2009; Willemsen *et al*., 2017). A Bayesian workflow in the open software environment R (R Core Team, 2022) via ‘brms’ (Bürkner, 2017) is illustrated for common PK studies including simple descriptive, and parallel designs with covariates, via two practical examples utilizing published data. Strategies for statistical inference and visualization of results are explored.

## Example 1: Application to Descriptive Pharmacokinetics

The classical theophylline dataset, originally reported by Upton and utilized by Beal et al., (2009) in early NONMEM work, and by Pinheiro and Bates (1995), among many other methodologic applications (Comets, Lavenu and Lavielle, 2017), was obtained from R (R Core Team, 2022). The data comprise observations of the plasma concentration of theophylline from 12 subjects over 24 hours, following oral administration. For the current work these data were selected as an archetypal example of a descriptive PK study with intensive sampling. For this application, the concentrations reported as zero were removed from the dataset; later, censoring will be implemented using HGAM, but in this dataset the censoring limit is not reported.

This paper emphasises the practical application and interpretation of hierarchical generalized additive models to non-compartmental analyses, and does not discuss mathematical details. Many resources are available that describe mathematical and statistical principles of HGAM and applications, at various levels of complexity (Pedersen *et al*., 2019; Van Rij *et al*., 2019; Wood, 2020). A hierarchical generalized additive model for the theophylline concentration at any point in time after administration was specified as:

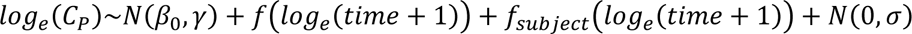

This model was expressed in R and ‘brms’ syntax as:

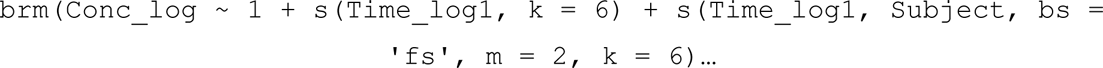

In which `Conc_log` is the log-transformed *C*_*P*_, `Time_log1` is the pseudo-log-transformed *time*, and `Subject` is a categorical variable indicating the subject. The model specification and notation here follows a highly recommended introduction to hierarchical GAM in R (Pedersen *et al*., 2019), based on the model syntax from package ‘mgcv’ (Wood, 2023). This specification denotes a population (‘global’) smooth effect of *time f*(), between-subject variation in the effect of *time* using the factor-smooth interaction term *f*_*subject*_(), between-subject variation (location shift) *γ* in the intercept *β*_0_, and normally-distributed constant conditional variation *σ*, all to predict the concentration *C*_*P*_ on *log*_*e*_ scale. The population smooth function in this case, where the basis is not specified, is a thin-plate regression spline (Wood, 2003). Note that the pseudo-logarithmic transformation of *time* allows *time* = 0 to be included (Jullion *et al*., 2009), but is somewhat non-linear especially for small values of *time*, which requires some caution to select an appropriate unit. This corresponds to a group-level model with shared wiggliness, as expressed in Pedersen et al., (2019). Analogous to the more general multilevel models, in which group-level parameters are drawn towards a common average, in this model the group-level functions are drawn towards a common average function, but with group-level variation in shape.

Parameters for this model include the smooth function variability for *time* and a coefficient for the linear trend of *time*, the intercept *β*_0_and its between-subject variation *γ*, and the conditional standard deviation σ. An important consideration is the specification of prior information for these parameters. Some parameters are readily interpretable. A prior distribution for the intercept represents the average *C*_*P*_ for average *time* (intercept priors are centered in ‘brms’), and *γ* is between-subject variation in this average, representing location-shift of the subjects. For this model these were *N*(0,5) for *β*_0_, intended to be weakly informative; *γ* is implemented within the smooth term. The prior distribution for σ was *HN*(0,1), which denotes that the for a sensible population pharmacokinetic model the scale of residual error, on logarithmic scale, should be reasonably small (less than 1). Priors for the smooth components are more difficult because they do not map to very interpretable statements about any real quantity, and each were retained using the package defaults; the linear trend of time was flat *U*(−∞, ∞), and the smooth function variance was *T*(0,2.5,3).

Estimation of the parameters was conducted via MCMC, by the No-U-Turn Sampler as implemented in Stan (Carpenter *et al*., 2017). Convergence and sufficiency of sampling were assessed using the *R*^ statistic (Vehtari *et al*., 2021) and effective sample sizes. Primary parameter estimates are reported in **Table 1**. The final model was strongly descriptive of the total variation in concentration (Bayes *R^2^*: 0.946, 90%CrI: 0.933, 0.957; Gelman et al., 2019). The fitted smooth functions, for each subject, are expressed visually (via ggplot2; Wickham, 2016) in **Figure 1**; overall the apparent fit of the model to the individual concentration data was adequate. There was no clear evidence of any major lack-of-fit for any subject. Residual analysis demonstrated that the constant-variation model was compromised by an apparent effect of time on residual variability (**Figure 2**), with residual variability declining with time. This pattern suggests relative difficulty in capturing the absorption phase.

**Figure 1:**
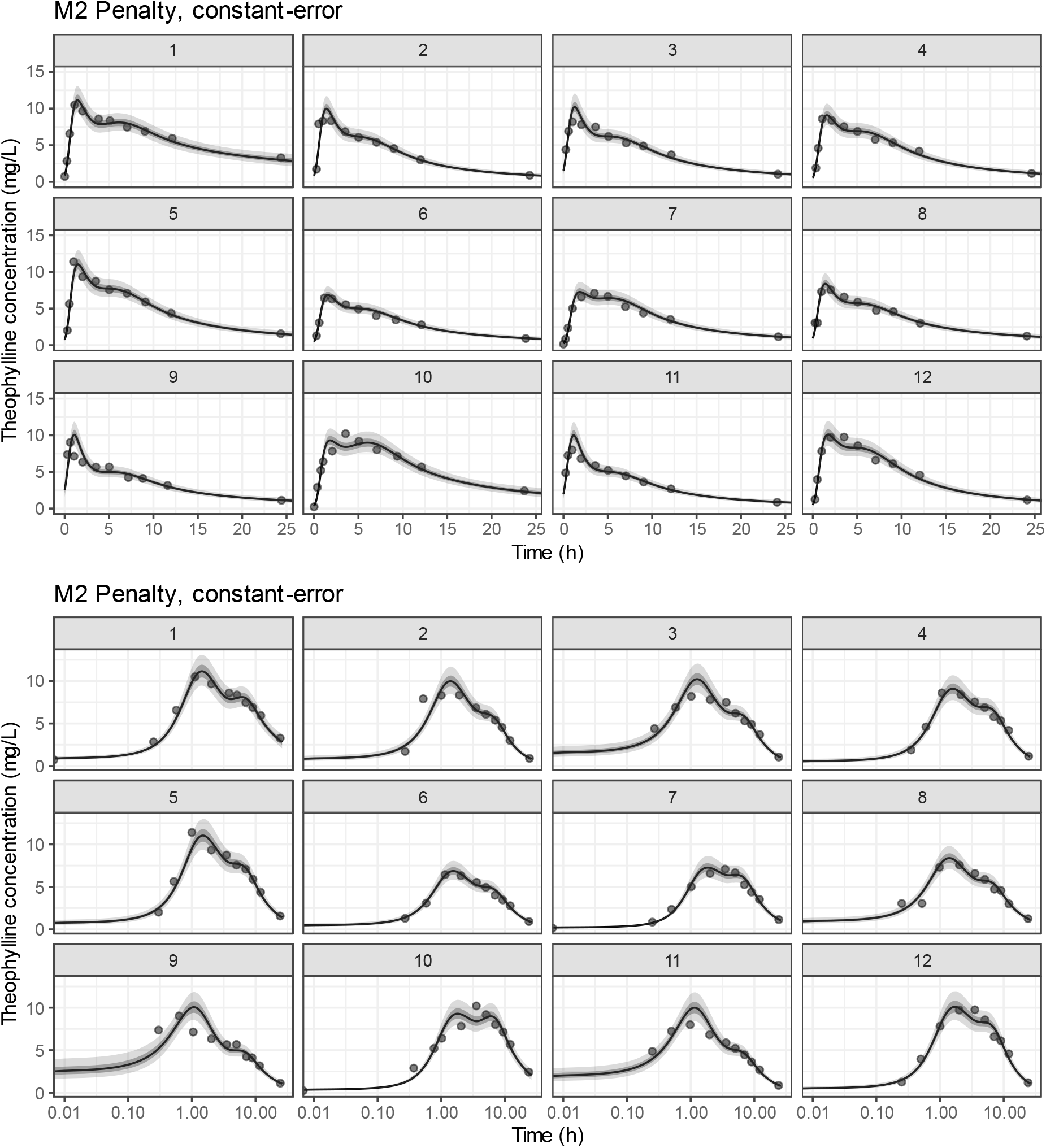
posterior-predicted theophylline concentrations over time, on linear or logarithmic time scale, conditional on subject. The line represents the posterior mean prediction, the shaded regions the 50% and 90% credible intervals, and the filled circles the observations of theophylline plasma concentration. Results are from the HGAM model for theophylline concentration over time, with constant residual error. The raw data were from the classical theophylline dataset reported by Upton, utilized by Beal et al., (2009) and Pinheiro and Bates (1995), and obtained here from R (R Core Team, 2022).

**Figure 2:**
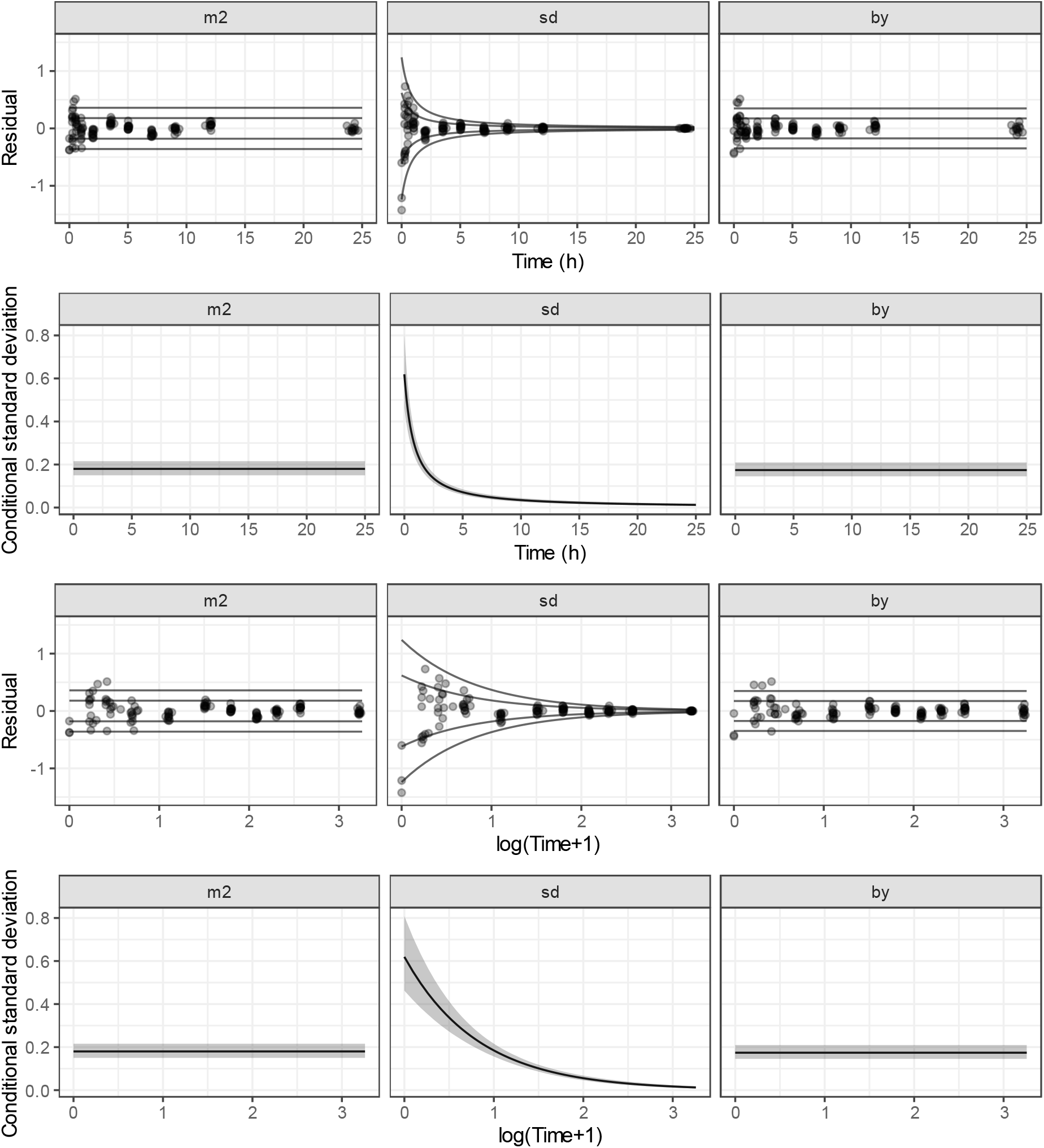
posterior mean residuals and conditional variation functions for the constant-error (‘m2’), time-varying error (‘sd’), and subject-varying wiggliness (‘by’) models, by time on either linear or pseudologarithmic scale. Solid lines in residual scatterplots represent ±1 and ±2 times (at the posterior mean) of the conditional variation. Solid line on the conditional standard deviation line plots represent the posterior mean of the conditional standard deviation on the log-scale, and the grey field its 90% credible interval. Results are from the HGAM models for theophylline concentration over time. The raw data were from the classical theophylline dataset reported by Upton, utilized by Beal et al., (2009) and Pinheiro and Bates (1995), and obtained here from R (R Core Team, 2022).

**Table 1:** posterior-predicted theophylline population *AUC*_*last*_, *T*_*MAX*_, and *C*_*MAX*_. Columns are the percentiles of the posterior distribution (credible limits and posterior median). m2: constant-error, constant-wiggliness model, sd: time-varying-error, constant-wiggliness model, by: varying wiggliness model. Results are from the HGAM model for theophylline concentration over time, with subject-varying wiggliness. The raw data were from the classical theophylline dataset reported by Upton, utilized by Beal et al., (2009) and Pinheiro and Bates (1995), and obtained here from R (R Core Team, 2022).

| Parameter | Model | 5% | 25% | 50% | 75% | 95% |
| --- | --- | --- | --- | --- | --- | --- |
| $AUC_{LAST}$ | m2 | 85.19 | 91.83 | 96.28 | 101.38 | 109.53 |
|  | sd | 87.44 | 93.43 | 97.39 | 101.48 | 107.85 |
|  | by | 88.28 | 96.55 | 102.74 | 108.76 | 118.14 |
| $T_{MAX}$ | m2 | 1.33 | 1.4 | 1.46 | 1.53 | 2.07 |
|  | sd | 1.69 | 1.8 | 1.89 | 1.99 | 2.15 |
|  | by | 1.37 | 1.44 | 1.5 | 1.56 | 6.13 |
| $C_{MAX}$ | m2 | 7.46 | 8.59 | 9.14 | 9.67 | 10.58 |
|  | sd | 7.6 | 8.17 | 8.59 | 9.03 | 9.71 |
|  | by | 7.62 | 9.43 | 10.04 | 10.62 | 11.51 |

A revised model was implemented:

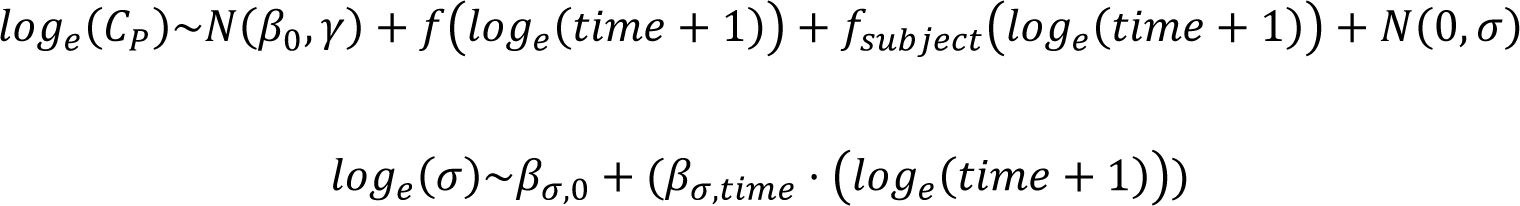

in R and ‘brms’ syntax as:

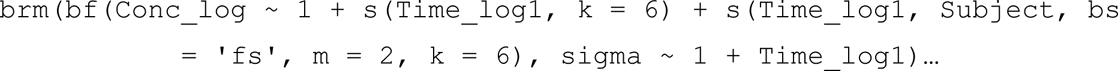

Regarding the expected value *C*_*P*_, the model is equivalent. Instead of constant conditional variation this model included a linear effect of time (on the *log*_*e*_ scale) on the conditional standard deviation σ. In this distributional model (Kneib, Silbersdorff and Säfken, 2023) both an intercept and linear coefficient for the effect of time on σ are estimated, on the *log*_*e*_ scale. Prior distributions were *N*(−1,1) for *β*_*σ*,0_ and *N*(0,2) for *β*_*σ*,*time*_, intended to denote that the expected σ is probably not larger than *e*^1^, and the effect of a 1-unit change in time on the *log*_*e*_ scale is most plausibly between -2 and 2 units; other priors were as for the constant variation model.

Overall the distributional model generated similar posterior predictions of the time-course of theophylline concentrations (**Figure 3**), to the constant-variation model (**Figure 1**). As the residual variation declined with time in this model, the fit is strongest for the later observations and weakest for the earliest observations, which may be reasonable given the greater complexity of the underlying biological processes earlier in time, but the scale of the conditional variation seems excessively small in this case. Residual analysis from the distributional model showed that the predicted conditional variation was not apparently contradicted by the observations (**Figure 2**). The proportion of described variation remained large (Bayes *R^2^:* 0.850, 90% CrI: 0.781, 0.904) but is more difficult to directly interpret under the non-constant variation. Though the time-varying model did improve the pattern of residuals, particularly around the early observations, the presence of time-varying error is difficult to explain. It would be surprising that analytic measurement error be time-varying; model lack-of-fit, or error in the exact sampling time, seem like more realistic possibilities.

**Figure 3:**
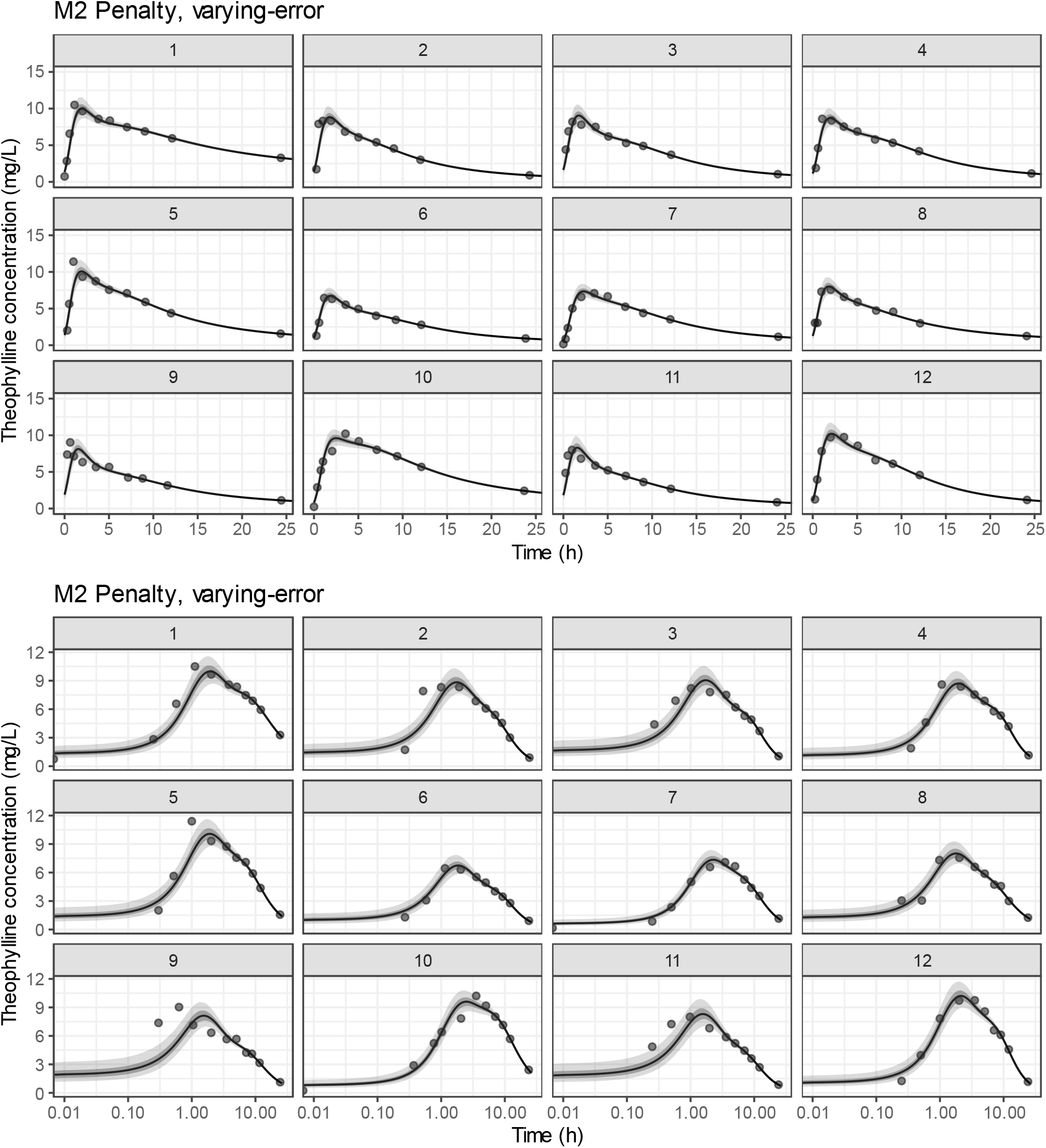
posterior-predicted theophylline concentrations over time, on linear or logarithmic time scale, conditional on subject. The line represents the posterior mean prediction, the shaded regions the 50% and 90% credible intervals, and the filled circles the observations of theophylline plasma concentration. Results are from the HGAM model for theophylline concentration over time, with time-varying conditional variation (residual error). The raw data were from the classical theophylline dataset reported by Upton, utilized by Beal et al., (2009) and Pinheiro and Bates (1995), and obtained here from R (R Core Team, 2022).

An additional modelling option is to allow greater flexibility for the form of the subject-level functions. A varying-wiggliness model (Pedersen *et al*., 2019) requires more parameters than the shared-wiggliness models, but may improve the fit for the individuals by reducing the degree of shrinkage. This model was specified as:

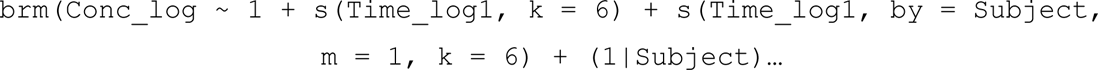

The time-course of the individual theophylline concentrations predicted by the model are shown in **Figure 4**, and are visually of better fit to the individual data than the shared-wiggliness models, particularly considering some of the more poorly-predicted observations (**Figure 1**). The proportion of variability in concentration explained was comparable (Bayes *R^2^*: 0.950, 90% CrI: 0.937, 0.960).

**Figure 4:**
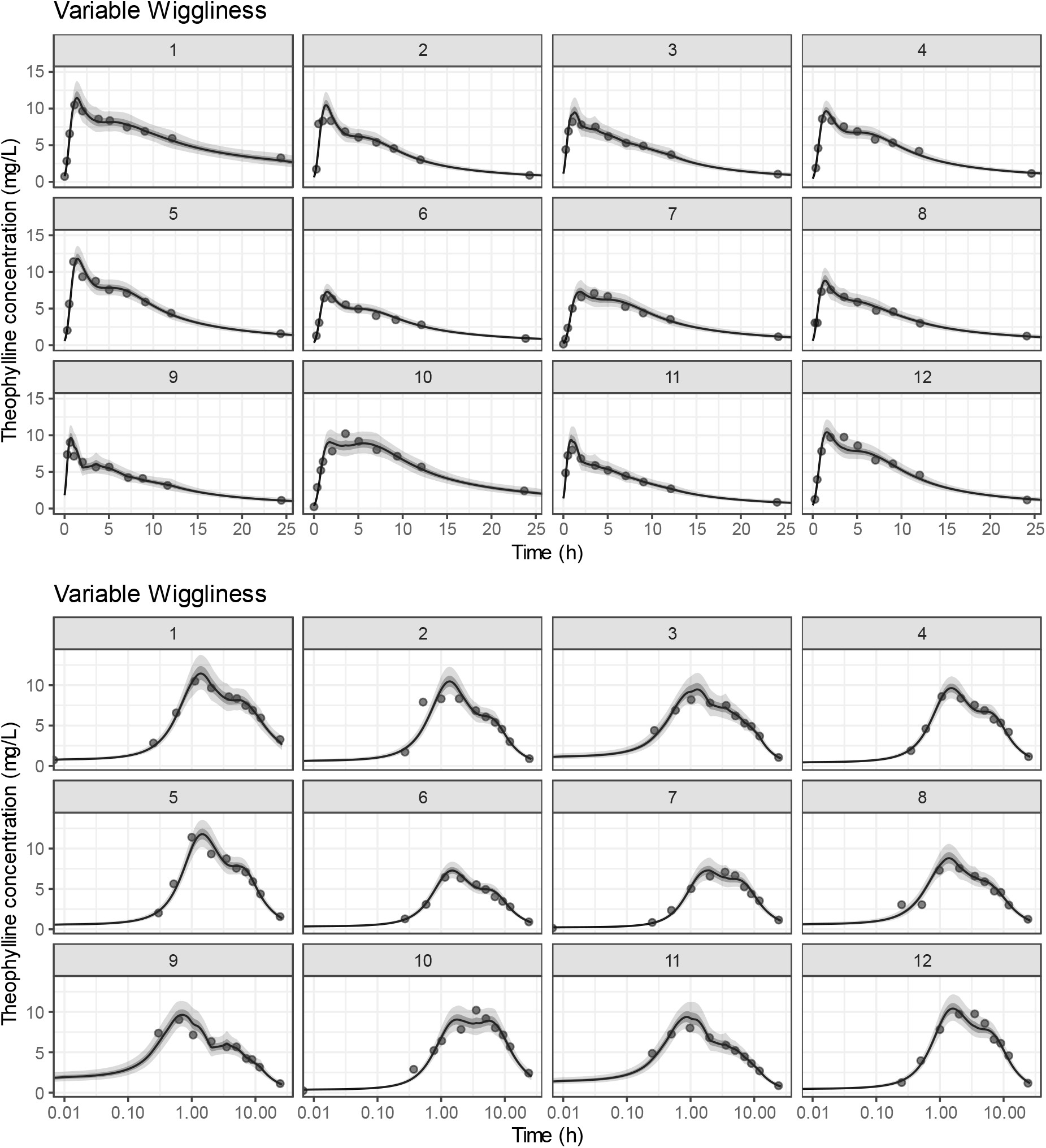
posterior-predicted theophylline concentrations over time, on linear or logarithmic time scale, conditional on subject. The line represents the posterior mean prediction, the shaded regions the 50% and 90% credible intervals, and the filled circles the observations of theophylline plasma concentration. Results are from the HGAM model for theophylline concentration over time, with subject-varying wiggliness. The raw data were from the classical theophylline dataset reported by Upton, utilized by Beal et al., (2009) and Pinheiro and Bates (1995), and obtained here from R (R Core Team, 2022).

Population predictions of the theophylline concentration from each model, conditional on an average subject (*i.e.* ignoring the subject-specific smooth terms) are expressed in **Figure 5**. The form of the population smooth functions was similar between the constant-error and varying-wiggliness models, but visibly different for the varying-error model. All were visually reasonable with respect to the aggregate data. Compared with classical graphics for pharmacokinetic data using sample statistics by nominal time, which do not represent a viable model for the expected concentration based on the data (Fossler, 2017), this graphic style using HGAM shows simultaneously the expected concentration for an average subject, its uncertainty, and the variability of the observed subjects.

**Figure 5:**
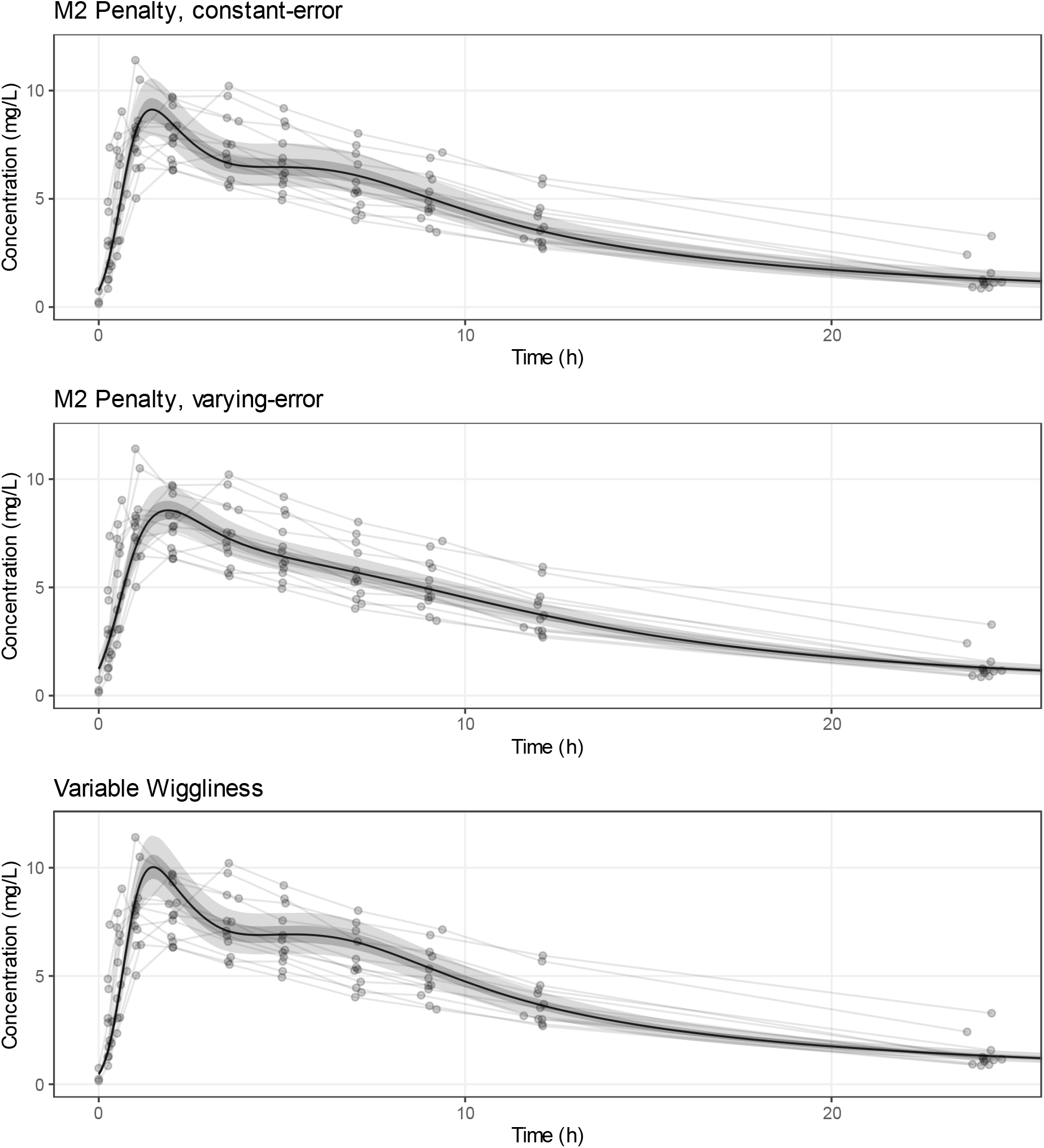
posterior-predicted population theophylline concentrations over time from the HGAM (various models by panel). The line represents the population posterior prediction (population smooth), the shaded regions the 50% and 90% credible intervals, the points the observations of theophylline concentration, and the grey lines join the observations by subject. This style of graphic is a possible replacement of the classic graphic style for a single-dose PK experiment, with the sample mean by nominal time as the central points, and the error bars its confidence interval, which is almost entirely uninformative (Fossler, 2017); no data is presented, and the confidence intervals for the sample means do not represent a sensible model. Results are from the HGAM model for theophylline concentration over time. The raw data were from the classical theophylline dataset reported by Upton, utilized by Beal et al., (2009) and Pinheiro and Bates (1995), and obtained here from R (R Core Team, 2022).

The key NCA parameter *AUC*_*last*_ was obtained via post-processing of the joint posteriors from each of the candidate models, by numeric integration on the interval 0-24h. Posterior distributions, both for the individuals and the population (a hypothetical average subject) were obtained by estimation from each sample. Because the per-sample numerical estimation is somewhat expensive this was fairly slow on a personal computer, taking several hours of computation while utilizing parallel processing by package ‘future’ (Bengtsson, 2021). For comparison, the *AUC*_*last*_ were generated by the trapezoidal method as implemented in the ‘ncappc’ package (Acharya *et al*., 2016).

The posterior distributions of population *AUC*_*last*_, from each model, are presented as histograms as implemented with ‘ggdist’ (Kay, 2022), in **Figure 6** along with the trapezoidal *AUC*_*last*_ and its confidence interval from percentile bootstrap via package ‘boot’ (Canty and Ripley, 2022). Summaries of the population posterior distributions for the *AUC*_*last*_, *C*_*MAX*_, and *T*_*MAX*_ are described in **Table 1**. The constant and time-varying error models generated similar posterior distributions, and intervals comparable to the bootstrap confidence intervals, while the posterior distributions from the varying-wiggliness model had greater width and skewness. As the subject-level estimates are drawn less strongly to the average in the varying-wiggliness model (less shrinkage), it is reasonable to expect that less information was available to support the population smooth, and its secondary parameters. Overall the posterior distributions for the individual *AUC*_*last*_ converged closely on the point estimates from the trapezoidal method (**Figures 7-9**), and posterior distributions were narrow, reflecting low, though not negligible, uncertainty.

**Figure 6:**
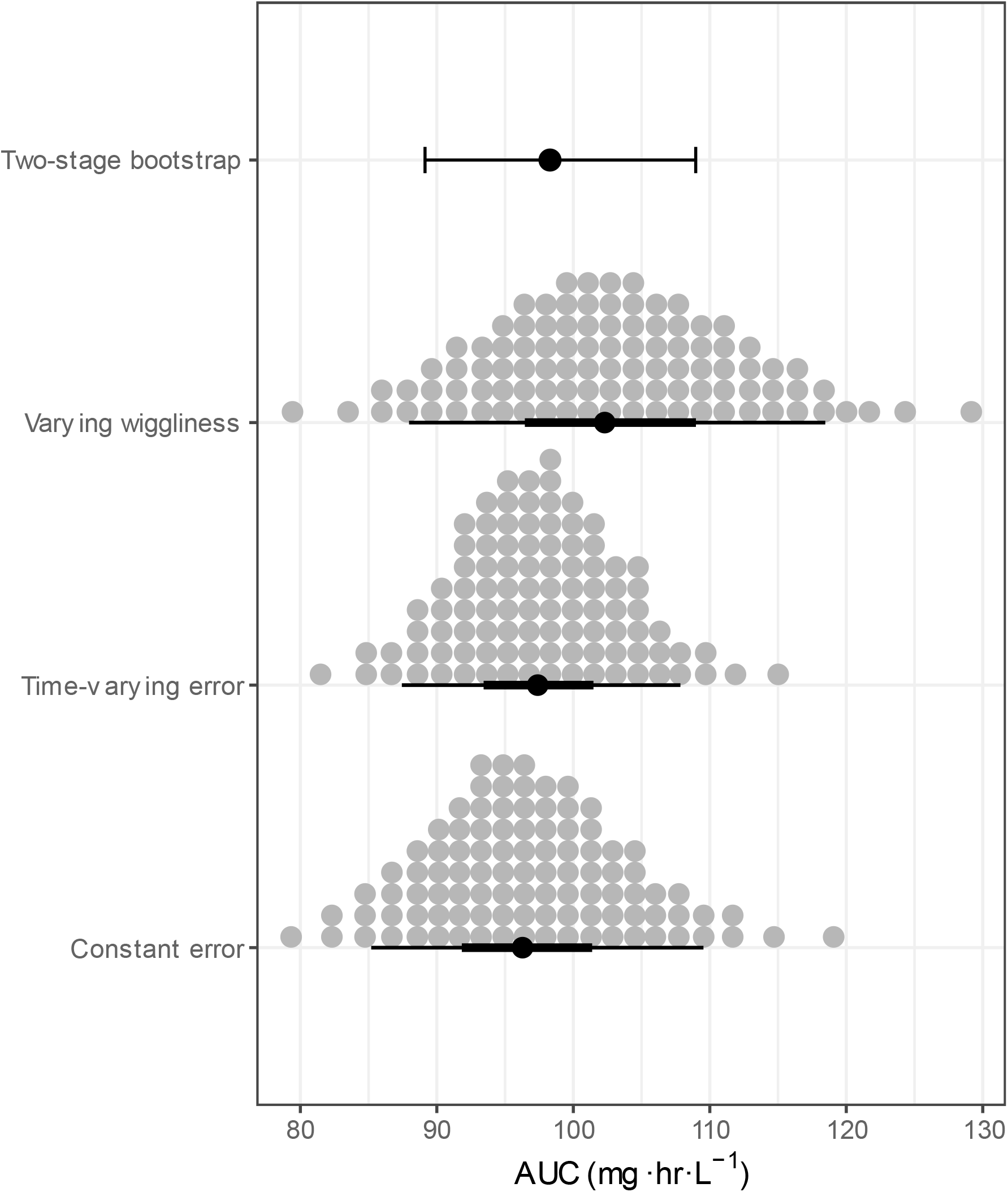
estimation of the theophylline population *AUC*_*last*_ under various models. Dot histograms (100 dots) represent posterior distributions of the theophylline *AUC*_*last*_ (from the global smoother), the solid points the posterior mean, and the bars the 50A% and 90% credible intervals. The solid point is the sample mean *AUC*_*last*_via the trapezoidal method, and the error bar its 90% confidence interval from percentile bootstrap. Results are from HGAM models for theophylline concentration over time. The raw data were from the classical theophylline dataset reported by Upton, utilized by Beal et al., (2009) and Pinheiro and Bates (1995), and obtained here from R (R Core Team, 2022).

**Figure 7:**
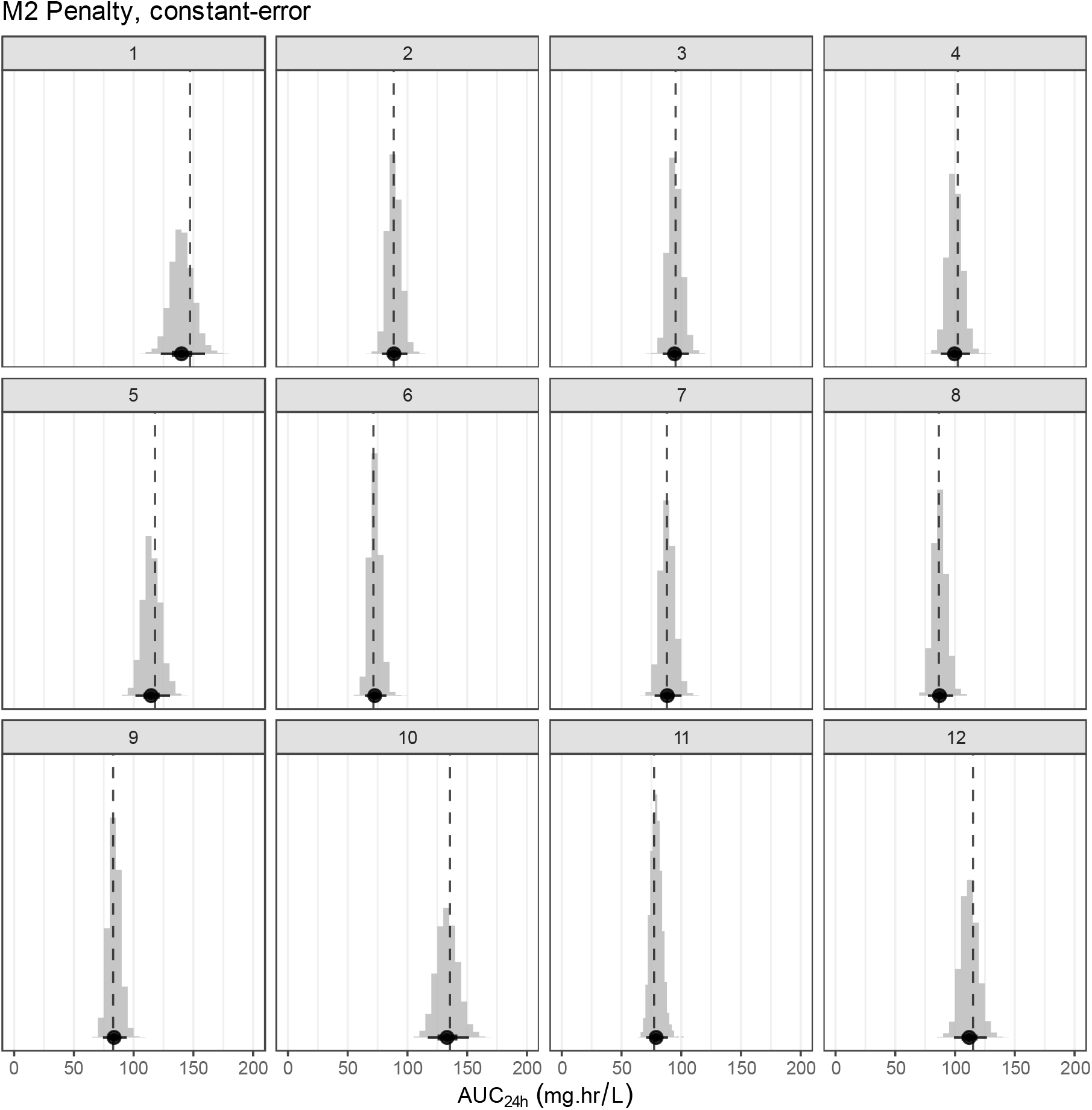
posterior distributions of the theophylline *AUC*_*last*_, conditional on subject. The posterior distribution is represented as the histogram. The point represents the posterior mean *AUC*_*last*_ and the bar the 90% credible interval. The dashed line is the corresponding point estimate from the trapezoidal method (conventional NCA). Results are from the HGAM model for theophylline concentration over time, with subject-varying wiggliness. The raw data were from the classical theophylline dataset reported by Upton, utilized by Beal et al., (2009) and Pinheiro and Bates (1995), and obtained here from R (R Core Team, 2022).

**Figure 8:**
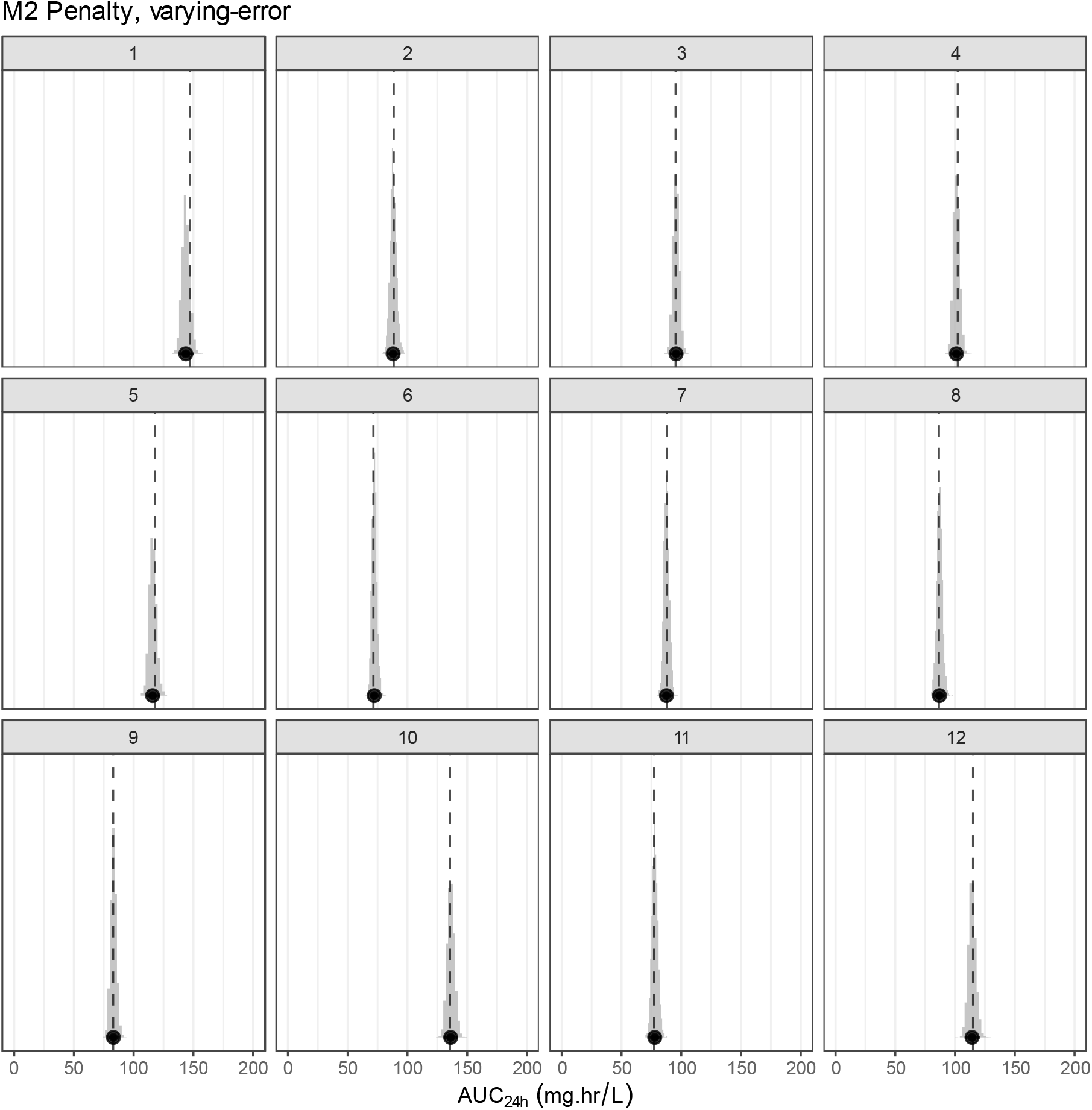
posterior distributions of the theophylline *AUC*_*last*_, conditional on subject. The posterior distribution is represented as the histogram. The point represents the posterior mean *AUC*_*last*_ and the bar the 90% credible interval. The dashed line is the corresponding point estimate from the trapezoidal method (conventional NCA). Results are from the HGAM model for theophylline concentration over time, with subject-varying wiggliness. The raw data were from the classical theophylline dataset reported by Upton, utilized by Beal et al., (2009) and Pinheiro and Bates (1995), and obtained here from R (R Core Team, 2022).

**Figure 9:**
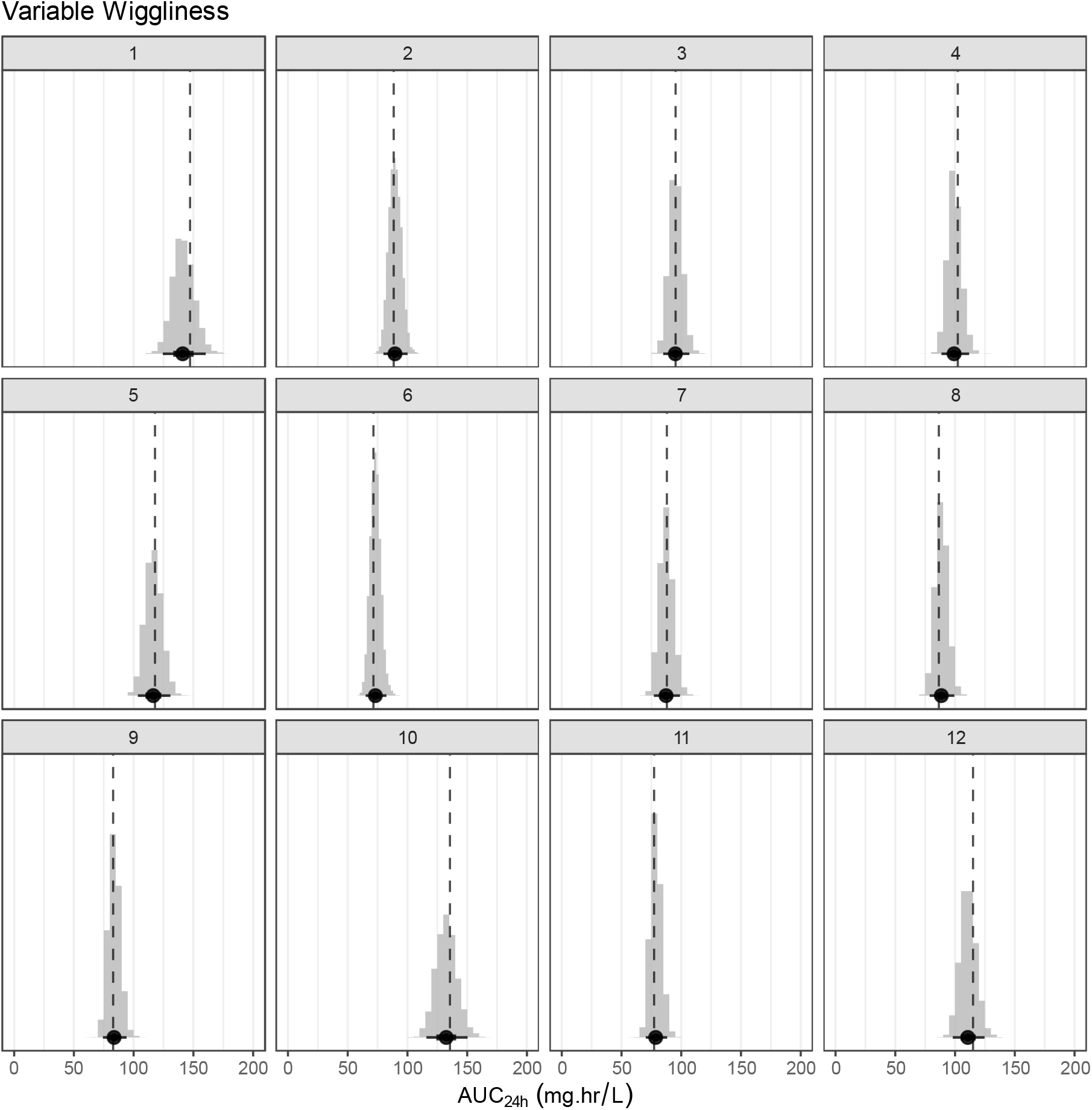
posterior distributions of the theophylline *AUC*_*last*_, conditional on subject. The posterior distribution is represented as the histogram. The point represents the posterior mean *AUC*_*last*_ and the bar the 90% credible interval. The dashed line is the corresponding point estimate from the trapezoidal method (conventional NCA). Results are from the HGAM model for theophylline concentration over time, with subject-varying wiggliness. The raw data were from the classical theophylline dataset reported by Upton, utilized by Beal et al., (2009) and Pinheiro and Bates (1995), and obtained here from R (R Core Team, 2022).

Noting the weak nature of the priors, especially for the smooth terms, the resulting posterior distributions for the pharmacokinetic parameters may not have strong interpretation in terms of belief, but rather represent mostly likelihood-based uncertainty statements. Compared to the various methods for confidence interval generation from NCA, mostly based on bootstrapping (Jaki, Wolfsegger and Ploner, 2009) which attempt to characterize uncertainty in a population parameter, this approach generates uncertainty statements corresponding to individual parameters. Because these statements are generated from a joint probability distribution for all relevant parameters, a credible region about *C*_*P*_, and a dependent credible interval about *AUC*_*last*_, which result from simultaneous uncertainty in multiple primary parameters, are relatively easy (though computationally expensive) to obtain. Similar statements may be available via the penalized likelihood approach in package ‘mgcv’. This contrasts with the relatively onerous procedures required for valid simultaneous confidence (*i.e.* frequentist) statements from population pharmacokinetic (i.e. nonlinear multilevel) models (Kümmel *et al*., 2018); notwithstanding the arguably poorer utility and interpretability of frequentist uncertainty statements (Morey *et al*., 2016).

Some computational challenges were noted during development of this model. Despite careful selection of the prior distributions, a small step size (`adapt_delta`) was necessary to avoid divergences (Betancourt, 2020), which are a threat to the validity of the results. In the author’s experience this is generally necessary for hierarchical additive models in ‘brms’, perhaps due to complex posterior geometry. A `max_treedepth` larger than the default was also required, which reflects sampling inefficiency (Stan Development Team, 2022); this may be resolvable using alternate parameterization, but none were clearly available for this case to the author’s knowledge.

## Example 2: Implementing Covariates

A popular workflow involving NCA is a ‘two-stage’ approach. Given some observed concentration-time data, the first stage is to estimate pharmacokinetic parameters for each subject-occasion by NCA. These parameter estimates (especially AUC) are then passed as data to a second stage, generally some linear model. Bioequivalence and bioavailability studies are a typical application. Notably, the first stage does not have to be NCA. Compartmental modelling by non-linear regression could be used instead, in which case the transition to a one-stage (‘population’) model could be implemented using a parametric, non-linear multilevel model as currently popular in pharmacometrics (Mould and Upton, 2012).

The hierarchical GAM suggests the possibility of an easy-to-implement, multilevel, ‘one-stage’ approach utilizing NCA, analogously to the relationship between nonlinear regression and nonlinear multilevel models. The theophylline model above, though emphasizing the conditional estimates of parameters for each subject, is a multilevel model; the subjects are variants from an estimated population effect and their estimates are drawn to it by partial pooling. Next, we will pursue an analysis with more explicit parameter estimation, including covariates, to illustrate some capabilities of a multilevel approach.

Warner and colleagues described the pharmacokinetics of meloxicam in dairy cattle after intravenous or oral administration (Warner et al., 2020b). The cows were at either mid-lactation or post-partum at the time of the study. Data were obtained from the repository (Warner et al., 2020a). From these data, a HGAM can be defined to estimate the effects of lactation stage *stage* and route of administration *route* on meloxicam concentration over time. The model was be specified as:

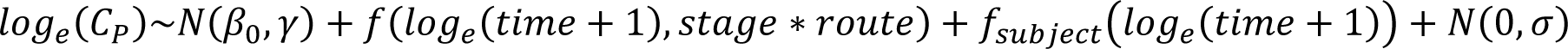

The specification of this model in R and ‘brms’ syntax was:

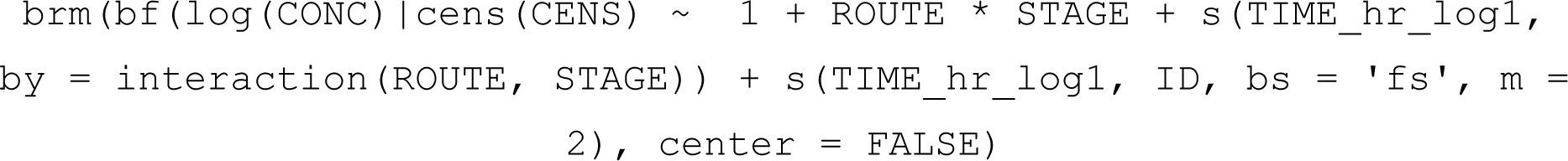

The terms have a similar meaning to the theophylline model above using the shared-wiggliness, constant-error structure. The additional *stage* ∗ *route* term denotes, as in the linear model setting, an interaction of the *stage* and *route* predictors; as well as the additive effect of each, the combination of both variables has an additional unique effect. In other words, there is a distinct population smooth predicted for each of the 4 combinations of the 2 binary predictors, and the predicted concentration-time relationship for some subject is implemented as the sum of one of these population smooths and a subject-specific smooth, where the subject-specific smooths vary commonly across categories.

Priors for the model were defined as in the theophylline model. An important note on the priors for this model is that they contain no information that reflects an expectation that the bioavailability of the oral route is lower than that of the intravenous route. Information about this feature must be obtained entirely from the data. Of course, no such prior information is available in any frequentist approach to this question (the majority of pharmacokinetic studies), so this is not ultimately a loss, except in the sense of the lost opportunity to provide additional information to the analysis. In the case where strong prior information is desired, fully-parametric pharmacokinetic models are available in which parameters have direct interpretation (Wakefield, 1996; Margossian, Zhang and Gillespie, 2022).

The presence of censoring in the concentration data is a complicating issue (Woodward and Whittem, 2019). The data contain numerous left-censored observations which represent meloxicam concentration below the lower performance limit of the analytic method. Fortunately, it is relatively simple to define these observations as left-censored in the model specification using the `cens` term (Bürkner, 2017). This implements the censoring in the likelihood function (Stan Development Team, 2022) such that the likelihood contribution from the censored observation is equal for any value below the censoring limit (representing that any value below the limit should be considered equally plausible), comparable to the method highlighted by Beal (2001).

A feature of this dataset is that the absorption phase was poorly observed; the earliest observations have concentrations well above the censoring limit. This initial phase is likely to be quite impactful on the pharmacokinetic parameter estimates. In the absence of a pre-dosing observation, the model is not constrained to any particular prediction at time zero, which may generate problematic predictions under the experimental assumption that drug is absent at time zero. The censoring model presents an opportunity to resolve this. By adding a left-censored observation at time zero, the model will be forced to conform to the *a priori* expected function shape with a starting point somewhere below the smallest drug concentration that could be detected. This synthetic data behaves essentially as prior information that expresses the knowledge (or the assumption) that drug concentration was below the censoring limit before drug was administered. This seems analogous to catalytic priors (Huang *et al*., 2022), in which the synthetic data is drawn from predictions by a simpler model. Of course, in cases where time zero observations were made, those observations can be utilized in the same way.

The estimation of parameters followed the same procedure as the theophylline model. No particular complications were encountered during sampling except for the requirement for a small step size to avoid divergent transitions. The conditional concentration-time relationship for each subject, highlighting their covariates, is expressed in **Figure 10**. The shape of the predicted function differed clearly between the IV and PO data, with the synthetic time zero observations apparently successful in generating the *a-priori* expected form of the absorption phase. Overall the model appeared to have reasonably low uncertainty in the conditional concentration-time function for each subject, though some apparently extreme observations were poorly fitted, reflecting the influence of shrinkage (**Figure 10)**.

**Figure 10:**
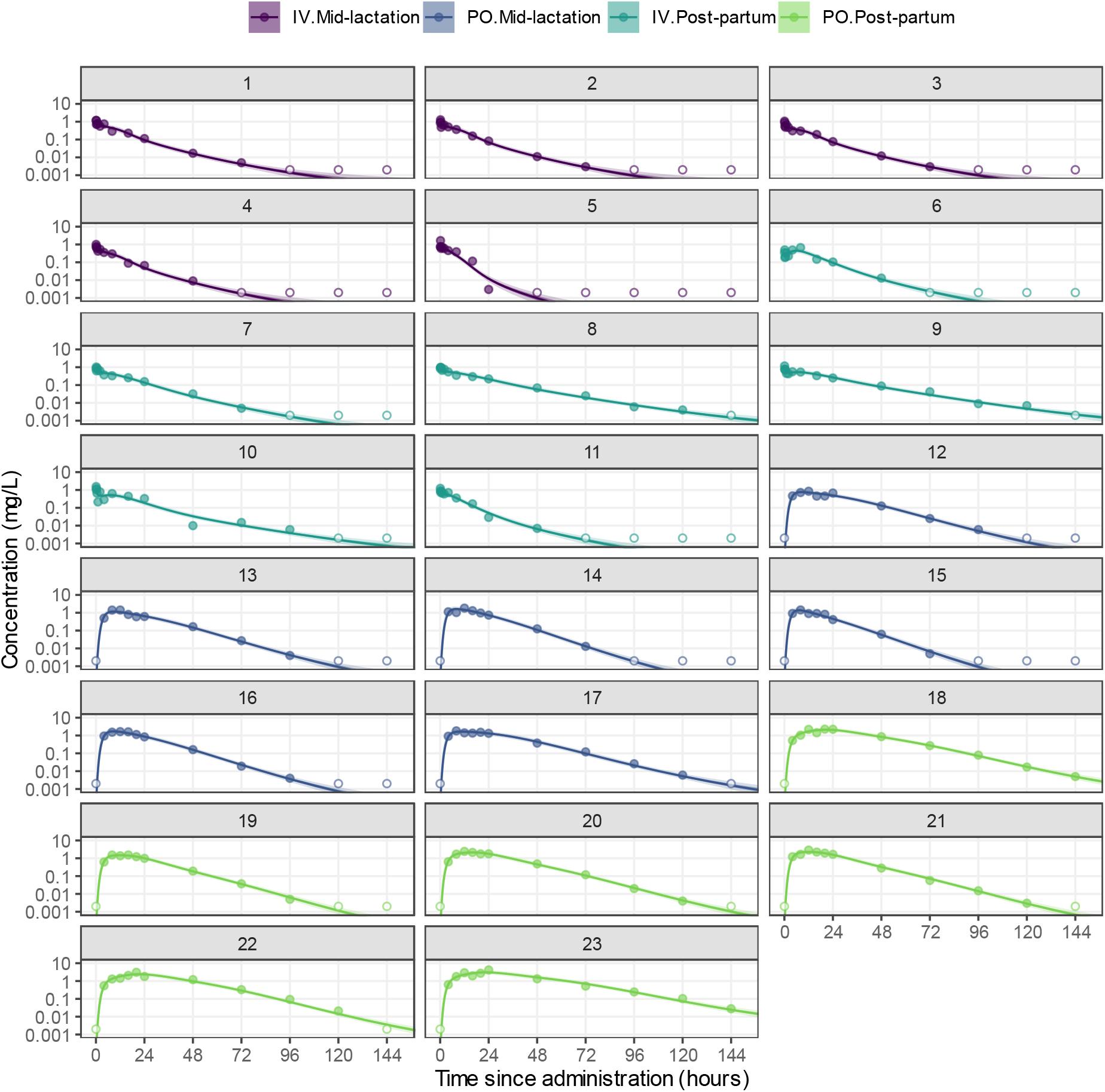
posterior-predicted meloxicam concentrations over time, conditional on subject, for intravenous or oral administration, in mid-lactation or post-partum cows. The line represents the posterior mean prediction, the shaded region the 90% credible intervals (which are narrow and sit close to the line on this scale), the filled circles the observations of meloxicam plasma concentration, and the open circles the left-censored observations (concentration below the analytic limit of quantification). Colors represent the covariates for route and stage effects. Note that the observations for PO at time 0 are synthetic and intended to express a prior belief that the concentration at time zero was too low, in principle, to be measured. Results are from the HGAM model for meloxicam concentration over time with covariates expressing route and stage effects. The raw data were obtained from Warner et al., (2020a).

Visualization of the ‘population’ smooths, conditional on the subject-level smooths set to zero, was utilized to highlight the covariate effects. These population smooths represent the model’s estimate of the underlying ‘average’ concentration-time relationship, by covariate condition. Those estimates, with credible regions, are expressed in **Figure 11**. The graphic shows a distinct population smooth function for each combination of the covariates, and a reasonable overall fit to the complete data. As for the theophylline example, this graphic style is a clean way to represent both the observed data and the estimated population effects, without the compromises inherent in traditional presentations (Fossler, 2017).

**Figure 11:**
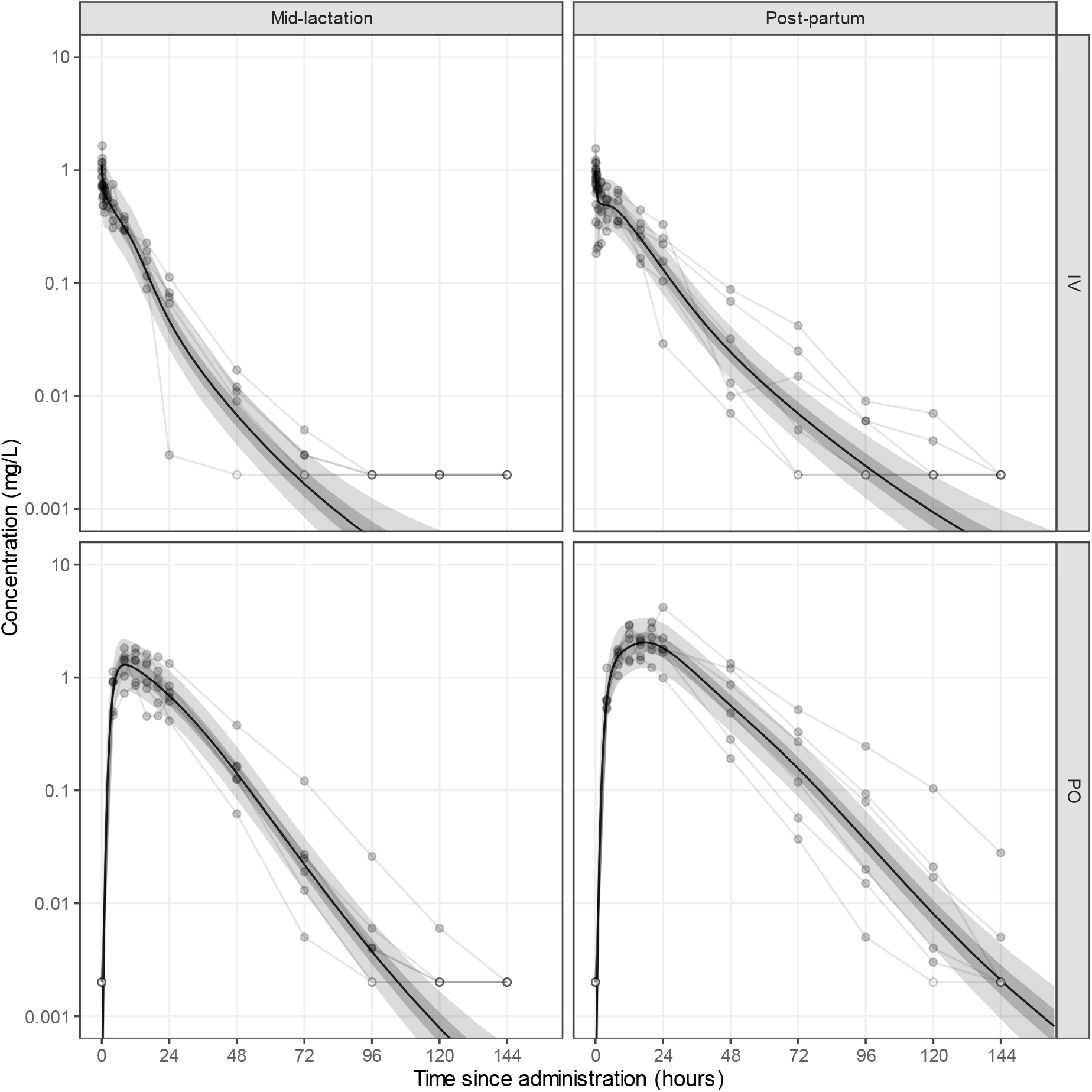
posterior-predicted population meloxicam concentrations over time (conditional on an ‘average’ subject), for intravenous or oral administration, in mid-lactation or post-partum cows, with concentration on logarithmic scale. The line represents the population posterior mean prediction, the shaded regions the 50% and 90% credible intervals, the filled circles the observations of meloxicam plasma concentration, and the open circles the left-censored observations (concentration below the analytic limit of quantification). The grey lines join the observations by subject. Note that the observations for PO at time 0 are synthetic and intended to express a prior belief that the concentration at time zero was too low, in principle, to be measured. Results are from the HGAM model for meloxicam concentration over time with covariates expressing route and stage effects. The raw data were obtained from Warner et al., (2020a).

A clear objective for the analysis is to express the covariate effects on the pharmacokinetic parameters. As for the theophylline model, the posterior distributions for the individual (conditional on subject) *AUC*_*last*_ (from 0 to 144h) were visualized as histograms (**Figure 12**), highlighting the covariates. Using posterior predictions from the population smooths, conditional on subject-level smooths set to zero, the covariate effects were expressed as contrasts in predicted *AUC*_*last*_. The Bayesian approach readily facilitates generation of such contrasts, via working directly with the samples from the posterior distribution. The approximate *AUC*_*last*_ was obtained by numeric integration of the predicted concentration-time function, for each posterior sample, for each condition (taking several hours of computation). Comparisons were generated of the *AUC*_*last*_ between stage for each of IV and PO administration *route*, and between route for each of mid-lactation and post-partum *stage*. Comparisons of *AUC*_*last*_ were generated between route, for PO administration. The resulting posterior distributions for *AUC*_*last*_ by condition, and for selected *AUC*_*last*_ ratios (contrasts) by condition, are expressed in **figure 13**.

**Figure 12:**
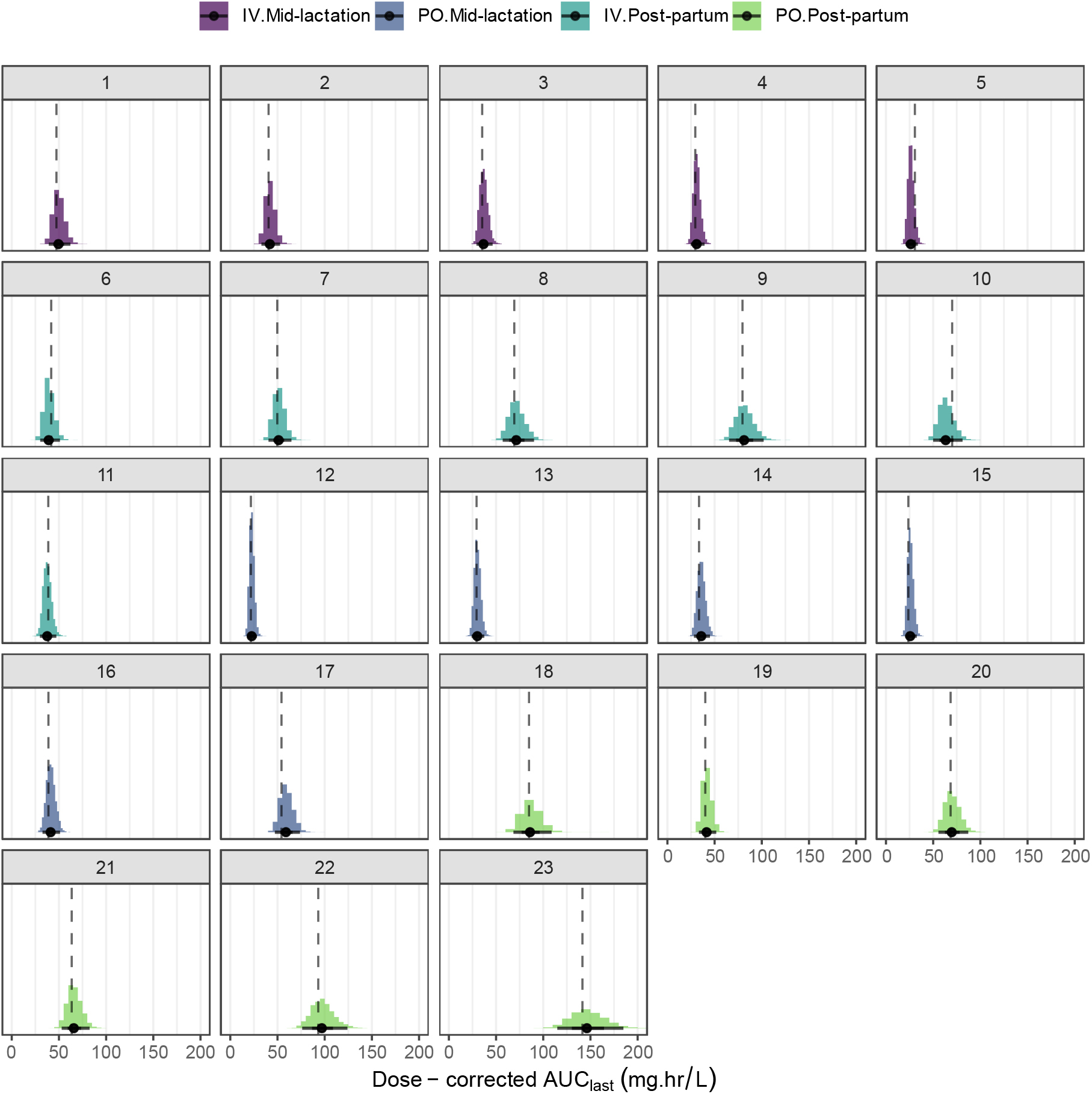
posterior distributions of the meloxicam *AUC*_*LAST*_ (dose-corrected to 1mg/kg), conditional on subject. The posterior distribution is represented as the histogram. The point represents the posterior mean *AUC*_*LAST*_ and the bar the 90% credible interval. The dashed line is the corresponding point estimate from the trapezoidal method (conventional NCA), in which censored observations were dropped. Colors represent the covariates for route and stage effects. Results are from the HGAM model for meloxicam concentration over time with covariates expressing route and stage effects. The raw data were obtained from Warner et al., (2020a).

**Figure 13:**
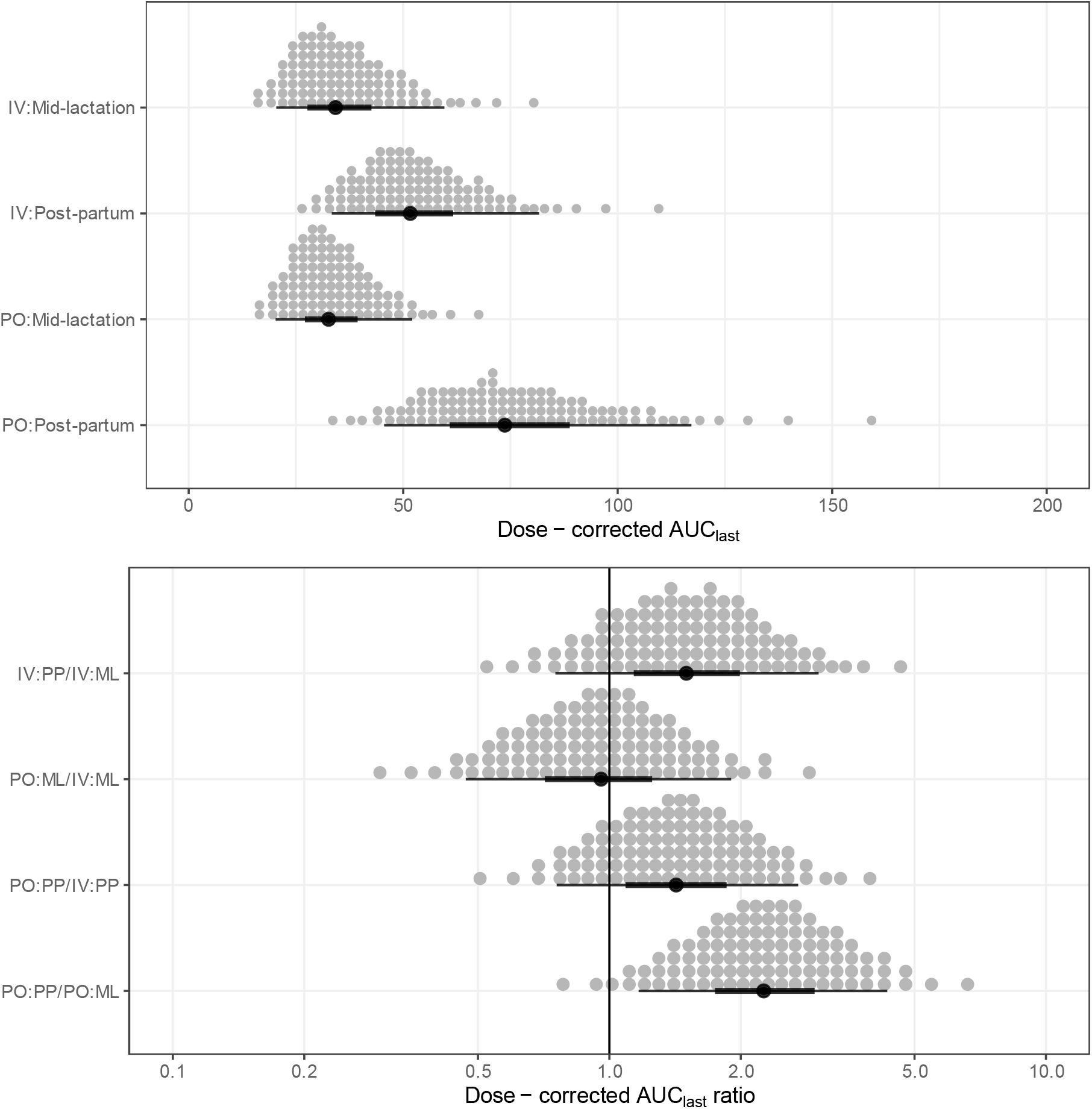
in the upper panel, posterior distributions, as 100 dots, of the population-predicted (conditional on a hypothetical ‘average’ subject, *i.e.* ignoring the between-subject variation), dose-corrected (*AUC*_*last*_:dose), for intravenous or oral administration, in mid-lactation or post-partum cows. In the lower panel, the population-predicted *AUC*_*last*_ ratio (relative exposure), for intravenous or oral administration, in mid-lactation or post-partum cows; the solid line represented *AUC*_*last*_ ratio of 1 (no effect). The points and intervals underneath the distributions represent the posterior mean, 50% and 90% credible intervals (posterior quantiles). Results are from the HGAM model for meloxicam concentration over time with covariates expressing dose and stage effects. The raw data were obtained from Warner et al., (2020a).

For comparison, a two-stage approach using a conventional NCA as implemented in the ‘ncappc’ package (Acharya *et al*., 2016), and a linear model, was applied. Censored observations were dropped. The resulting linear predictions of the geometric mean *AUC*_*last*_ by condition, and their ratios, were generated by percentile bootstrap as implemented in package ‘boot’ (Canty and Ripley, 2022) along with confidence intervals. Notably, these uncertainty statements do not correspond closely to the population predictions from the HGAM model. The HGAM population estimates correspond to the estimate of the *AUC*_*last*_ of underlying population function, *i.e.* the expected *AUC*_*last*_ for a hypothetical average subject, whereas the two-stage estimates correspond to the geometric mean of the subjects’ *AUC*_*last*_, *i.e.* the observed subjects’ *AUC*_*last*_ on average. A comparable statement from the HGAM can be generated easily, after some computation; the posterior distribution of the conditional *AUC*_*last*_ for each subject was generated, and the geometric mean and ratios for those *AUC*_*last*_ result in posterior distributions for those parameters. These are expressed in **figure 14**. Summarized as point estimates and intervals, these are of comparable value to the two-stage bootstrap results (noting of course that these credible intervals are not equivalent to confidence intervals; Morey et al., 2016), and both are much narrower than the population estimate from the HGAM.

**Figure 14:**
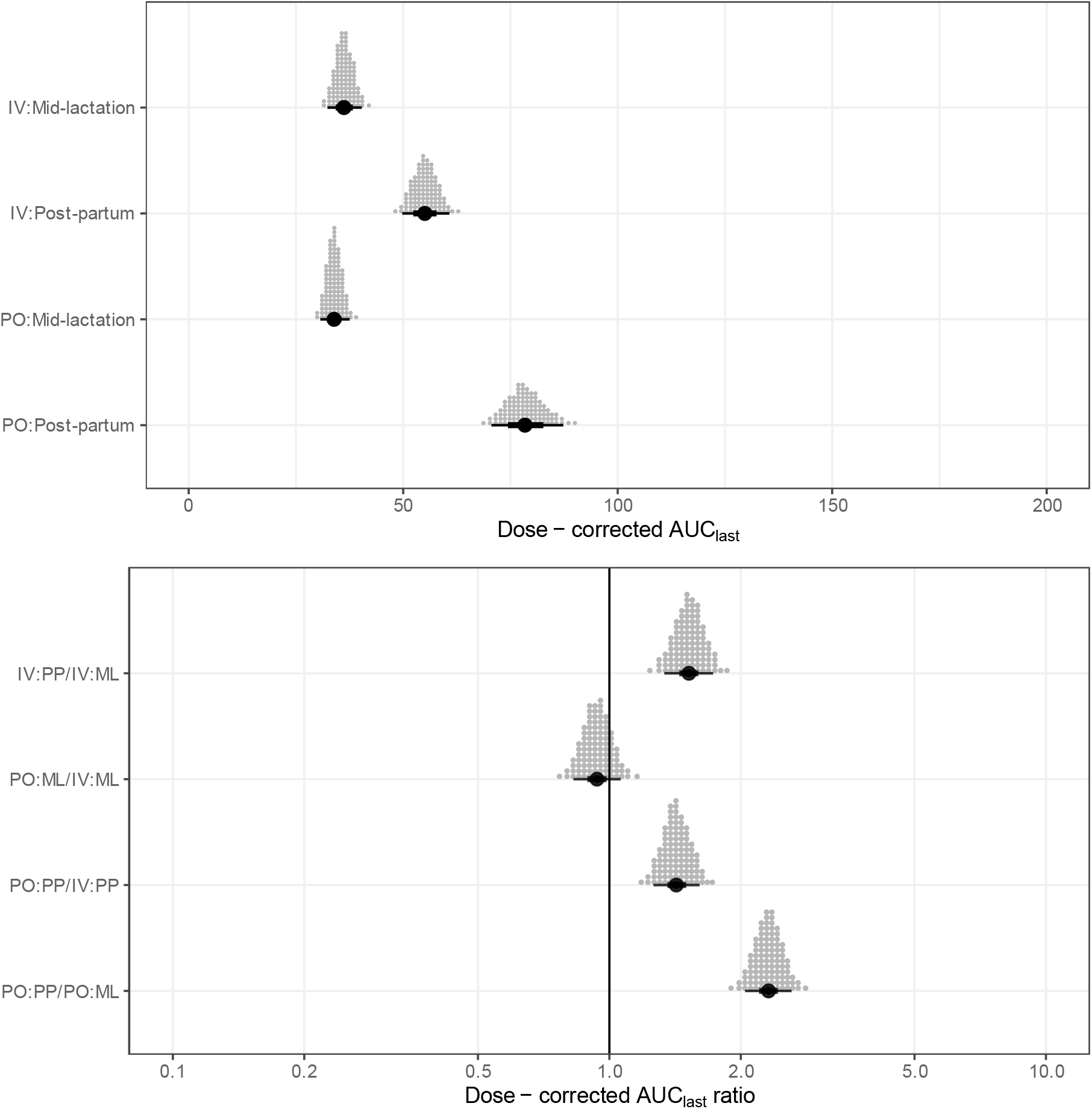
in the upper panel, posterior distributions, as 100 dots, of the predicted mean *AUC*_*last*_, conditional on the observed subjects (*i.e.* estimated average of the studied subjects), dose-corrected (*AUC*_*last*_:dose), for intravenous or oral administration, in mid-lactation or post-partum cows. In the lower panel, the predicted mean *AUC*_*last*_ratio (relative exposure) conditional on the observed subjects, for intravenous or oral administration, in mid-lactaion or post-partum cows; the solid line represented *AUC*_*last*_ ratio of 1 (no effect). The points and intervals underneath the distributions represent the posterior mean, 50% and 90% credible intervals (posterior quantiles). Results are from the HGAM model for meloxicam concentration over time with covariates expressing dose and stage effects. The raw data were obtained from Warner et al., (2020a).

## Discussion

These explorations suggest potential utility of the HGAM approach for pharmacokinetic data analysis. Bringing the partial pooling, likelihood-based parameter estimation, and uncertainty quantification that are generally features of population pharmacokinetics to NCA, the method facilitates a more modern, single-stage statistical approach utilizing all the available information. In these examples, the estimation of parameters was apparently accurate in comparison to a conventional workflow. These models were constructed via a Bayesian approach, allowing posterior distributions for functions of the model, such as areas or maxima, to be easily obtained as functions of the samples. As the prior information is overall quite weak, especially regarding the parameters of the smooth functions which lack a clear interpretation (and were simply left at the package defaults), the resulting posteriors may lack a strong interpretation in terms of belief. However, as a pragmatic choice this may be reasonable, if the model is computationally efficient enough.

Examples focussed on the analysis of multiple-subject data, analogously to the typical workflow for nonlinear multilevel modelling. Though it is possible to simply generate a separate GAM for each subject, in principle allowing for some features such as uncertainty quantification and censoring, this is not likely to be an efficient usage of the data in most instances, so was not characterized in detail here. Similarly possible is the implementation of HGAM as the first stage in an otherwise two-stage approach, perhaps for evaluation of covariates, in which case the potential advantages compared to the trapezoidal method include partial pooling, implementation of censoring, and uncertainty quantification. This may be particularly advantageous in sparse-data situations, if NCA is desired rather than a parametric model. Additionally, uncertainty in the pharmacokinetic parameter estimates, expressed for example as the posterior standard deviation, could be transferred to the second stage via a measurement error model, or by multiple imputation. Though joint models would more readily facilitate downstream usage of the parameter estimates from this model in some other estimation, to my knowledge this is currently technically limited by the pre-processing generation of smooth terms in ‘brms’ which is not easily accessible in custom Stan programs, precluding definitions of smooths which depend on parameters. Future developments of HGAM directly in Stan would facilitate such applications. Of course, spline models such as penalized splines may be directly implemented by the user in Bayesian analysis software, as by Jullion et al., (2009), but this undermines accessibility for non-mathematical users.

Interesting extensions to the model could be considered in future. The thin-plate regression spline (Wood, 2003) as applied for these analyses is capable of modelling interactions of smooths; this could, for example, be used to implement an effect of dose, in cases where the influence of dose variability on the concentration-time relationship is required or of interest. The form of the dose-exposure relationship may present a challenge in this case, suggesting the need for careful selection of priors or spline degrees of freedom. A key application for NCA is the evaluation of bioequivalence (BE). Similarly to the MSITAR approach explored by Willemsen et al., (2017), the HGAM approach may be suitable for BE studies, where formulation or route are implemented as predictors, similarly to the meloxicam example. For more complex BE designs, with crossover for example, extensions of this model for additional levels to implement within-subject varying splines would be required. Such models could be well-suited to BE statements, either conditional on the tested subjects or conditional on the population smooth, potentially acting as an intermediate case, between the two-stage approach using typical NCA and a population pharmacokinetic approach with a fully-parametric model (Dubois *et al*., 2011). Population BE statements marginalizing the between-subject variation as suggested by (Willemsen *et al*., 2017), from ‘brms’ models are a more complex consideration, especially regarding the potential cost of computation and the random-effects definition of the spline terms to conduct this marginalization in post-processing (Wiley and Hedeker, 2022). Estimation of model-predicted AUC under the current workflow is computationally expensive due to multiple calls to the ‘brms’ model during numeric integration, which could perhaps be improved using a trapezoidal method instead.

A specific limitation of the model warrants specific mention. These splines perform poorly in extrapolation beyond the observed data, simply because the form of the function no longer has any support. Though it is computationally feasible to simply generate *AUC*_∞_ by numeric integration from the fitted model just as for any other endpoint, in practice the improper integrals, conditional on subject, converged poorly across samples, generating sufficient divergent estimates in many cases that these posterior distributions were of suspect reliability. In typical NCA the extrapolation to infinity is achieved using a simple linear function on the log-scale; this was previously implemented in a spline approach to NCA (Jullion *et al*., 2009), but its validity for the current HGAM model is not clear. Future evaluations could consider shape-constrained additive models (SCAM) which present the possibility of enforcing monotone or concave splines for example, including for multiple predictors (Pya and Wood, 2015). Of course, in studies with suitably sensitive analytic methods and observations over a sufficient length of time, *AUC*_*last*_ approaches *AUC*_∞_, such that resulting bias in simply using *AUC*_*last*_ is likely to be small and generally irrelevant, though this is a matter for careful evaluation by the user. Though this study focussed on ‘primary’ parameters *AUC*, *T*_*MAX*_, and *C*_*MAX*_, some of the more complex NCA parameters (Veng-Pedersen, 1984), including those determined from multiple dosing events, would fit naturally in the multilevel approach as they could be jointly estimated, and could be considered in future applications.

In summary, the proposed HGAM approach to PK analysis overcomes many of the disadvantages of more traditional two-stage approaches for descriptive PK studies, but without the demands of model selection imposed by typical non-linear multilevel modelling. Compared to previous demonstrations of semi-parametric modelling for NCA (Park *et al*., 1997; Jullion *et al*., 2009; Willemsen *et al*., 2017), the HGAM approach appears to be at least as flexible, whilst also being supported by an accessible open-source implementation. Priorities for further evaluation of this model include application to sparse datasets, inclusion of dose variation, and extension to within-subject variation via additional grouping levels.

**Table 2:** posterior-predicted meloxicam *AUC*_*last*_ by STAGE and ROUTE conditions, for intravenous (IV) or oral (PO) administration, in mid-lactation (ML) or post-partum (PP) cows. Point estimates are the posterior median of the population *AUC*_*last*_**^A^,** posterior median of the geometric mean *AUC*_*last*_ across subjects**^B^,** or maximum likelihood estimate of the geometric mean *AUC*_*last*_**^C^**. Interval estimates are credible interval limits, and for the trapezoidal method, confidence limits from percentile bootstrap*. Results are from the HGAM model for meloxicam concentration over time with covariates expressing dose and stage effects. The raw data were obtained from Warner et al., (2020a).

| Model | Condition | 5% | 25% | Point Estimate | 75% | 95% |
| --- | --- | --- | --- | --- | --- | --- |
| HGAM, population prediction | IV:ML | 20.42 | 27.66 | 34.23<sup>A</sup> | 42.6 | 59.61 |
|  | PO:ML | 20.25 | 27.1 | 32.61<sup>A</sup> | 39.38 | 52.09 |
|  | IV:PP | 33.34 | 43.46 | 51.65<sup>A</sup> | 61.65 | 81.68 |
|  | PO:PP | 45.55 | 60.87 | 73.7<sup>A</sup> | 88.81 | 117.19 |
| HGAM, predicted geometric mean | IV:ML | 32.92 | 34.84 | 36.18<sup>B</sup> | 37.65 | 39.72 |
|  | PO:ML | 31.23 | 32.77 | 33.88<sup>B</sup> | 35.1 | 36.87 |
|  | IV:PP | 50.63 | 53.22 | 55.05<sup>B</sup> | 56.95 | 59.71 |
|  | PO:PP | 71.82 | 75.59 | 78.39<sup>B</sup> | 81.36 | 85.77 |
| Trapezoidal method, predicted mean* | IV:ML | 31.91 | 34.12 | 36.09<sup>C</sup> | 38.27 | 40.92 |
|  | PO:ML | 25.94 | 29.21 | 31.99<sup>C</sup> | 35.22 | 39.55 |
|  | IV:PP | 46.32 | 52.54 | 56.08<sup>C</sup> | 61.01 | 68.37 |
|  | PO:PP | 58.38 | 68.63 | 76.19<sup>C</sup> | 84.67 | 99.88 |

**Table 3:** posterior-predicted meloxicam *AUC*_*last*_ ratio (contrasts) by STAGE and ROUTE conditions, for intravenous (IV) or oral (PO) administration, in mid-lactation (ML) or post-partum (PP) cows. Point estimates are the posterior median of ratios of population *AUC*_*last*_**^A^,** posterior median of the geometric mean *AUC*_*last*_ across subjects**^B^,** or maximum likelihood estimate of the geometric mean *AUC*_*last*_**^C^**. Interval estimates are credible interval limits (posterior percentiles), and for the trapezoidal method, confidence limits from percentile bootstrap**^†^**. Results are from the HGAM model for meloxicam concentration over time with covariates expressing dose and stage effects. The raw data were obtained from Warner et al., (2020a).

| Model | Contrast | 5% | 25% | Point Estimate | 75% | 95% |
| --- | --- | --- | --- | --- | --- | --- |
| HGAM, population prediction | IV:PP/IV:ML | 0.75 | 1.14 | 1.5<sup>A</sup> | 1.99 | 3.01 |
|  | PO:PP/PO:ML | 1.17 | 1.75 | 2.26<sup>A</sup> | 2.95 | 4.33 |
|  | PO:ML/IV:ML | 0.47 | 0.71 | 0.96<sup>A</sup> | 1.25 | 1.9 |
|  | PO:PP/IV:PP | 0.76 | 1.09 | 1.42<sup>A</sup> | 1.86 | 2.71 |
| HGAM, predicted mean | IV:PP/IV:ML | 1.34 | 1.45 | 1.52<sup>B</sup> | 1.6 | 1.73 |
|  | PO:PP/PO:ML | 2.05 | 2.2 | 2.31<sup>B</sup> | 2.43 | 2.61 |
|  | PO:ML/IV:ML | 0.83 | 0.89 | 0.94<sup>B</sup> | 0.99 | 1.06 |
|  | PO:PP/IV:PP | 1.26 | 1.36 | 1.42<sup>B</sup> | 1.5 | 1.61 |
| Trapezoidal method, predicted mean<sup>†</sup> | IV:PP/IV:ML | 1.25 | 1.43 | 1.55<sup>C</sup> | 1.71 | 1.93 |
|  | PO:PP/PO:ML | 1.7 | 2.08 | 2.38<sup>C</sup> | 2.75 | 3.38 |
|  | PO:ML/IV:ML | 0.69 | 0.8 | 0.89<sup>C</sup> | 0.98 | 1.13 |
|  | PO:PP/IV:PP | 0.99 | 1.19 | 1.36<sup>C</sup> | 1.54 | 1.89 |

## References

Acharya, C. et al. (2016) ‘A diagnostic tool for population models using non-compartmental analysis: The ncappc package for R’, Computer Methods and Programs in Biomedicine, 127, pp. 83–93. Available at: https://doi.org/10.1016/j.cmpb.2016.01.013.

Barnett, H.Y. et al. (2021) ‘Methods for Non-Compartmental Pharmacokinetic Analysis With Observations Below the Limit of Quantification’, Statistics in Biopharmaceutical Research, 13(1), pp. 59–70. Available at: https://doi.org/10.1080/19466315.2019.1701546.

Beal, S.L. (2001) ‘Ways to Fit a PK Model with Some Data Below the Quantification Limit’, Journal of Pharmacokinetics and Pharmacodynamics, 28(5), pp. 481–504. Available at: https://doi.org/10.1023/A:1012299115260.

Beal, S.L. et al. (2009) NONMEM User’s Guides (1989-2009). Ellicott City, MD, USA: Icon Development Solutions.

Bengtsson, H. (2021) ‘A Unifying Framework for Parallel and Distributed Processing in R using Futures’, The R Journal, 13(2), p. 208. Available at: https://doi.org/10.32614/RJ-2021-048.

Betancourt, M. (2020) Hierarchical Modeling. Available at: https://github.com/betanalpha/knitr_case_studies/tree/master/hierarchical_modeling, commit 27c1d260e9ceca710465dc3b02f59f59b729ca43.

Bürkner, P.-C. (2017) ‘brms: An R Package for Bayesian Multilevel Models Using Stan’, Journal of Statistical Software, 80(1). Available at: https://doi.org/10.18637/jss.v080.i01.

Canty, A. and Ripley, B. (2022) ‘boot: Bootstrap R (S-Plus) Functions’.

Carpenter, B. et al. (2017) ‘*Stan*: A Probabilistic Programming Language’, Journal of Statistical Software, 76(1). Available at: https://doi.org/10.18637/jss.v076.i01.

Chiou, W.L. (1978) ‘Critical evaluation of the potential error in pharmacokinetic studies of using the linear trapezoidal rule method for the calculation of the area under the plasma level-time curve’, Journal of Pharmacokinetics and Biopharmaceutics, 6(6), pp. 539–546. Available at: https://doi.org/10.1007/BF01062108.

Comets, E., Lavenu, A. and Lavielle, M. (2017) ‘Parameter Estimation in Nonlinear Mixed Effect Models Using **saemix**, an *R* Implementation of the SAEM Algorithm’, Journal of Statistical Software, 80(3). Available at: https://doi.org/10.18637/jss.v080.i03.

Dominici, F. (2002) ‘On the Use of Generalized Additive Models in Time-Series Studies of Air Pollution and Health’, American Journal of Epidemiology, 156(3), pp. 193–203. Available at: https://doi.org/10.1093/aje/kwf062.

Dubois, A. et al. (2011) ‘Model-based analyses of bioequivalence crossover trials using the stochastic approximation expectation maximisation algorithm’, Statistics in Medicine, 30(21), pp. 2582–2600. Available at: https://doi.org/10.1002/sim.4286.

Dunne, A. and King, P. (1989) ‘Estimation of noncompartmental parameters: A technical note’, Journal of Pharmacokinetics and Biopharmaceutics, 17(1), pp. 131–137. Available at: https://doi.org/10.1007/BF01059092.

Fossler, M.J. (2017) ‘Some Thoughts About the Mean Concentration-Versus-Time Plot’, Clinical Pharmacology in Drug Development, 6(3), pp. 220–223. Available at: https://doi.org/10.1002/cpdd.353.

Gabrielsson, J. and Weiner, D. (2012) ‘Non-compartmental Analysis’, in B. Reisfeld and A.N. Mayeno (eds) Computational Toxicology: Volume I. Totowa, NJ: Humana Press, pp. 377–389. Available at: https://doi.org/10.1007/978-1-62703-050-2_16.

Gelman, A. et al. (2019) ‘R-squared for Bayesian Regression Models’, The American Statistician, 73(3), pp. 307–309. Available at: https://doi.org/10.1080/00031305.2018.1549100.

Gelman, A., Bois, F. and Jiang, J. (1996) ‘Physiological Pharmacokinetic Analysis Using Population Modeling and Informative Prior Distributions’, Journal of the American Statistical Association, 91(436), pp. 1400–1412. Available at: https://doi.org/10.1080/01621459.1996.10476708.

Greenland, S. (2000) ‘Principles of multilevel modelling’, International Journal of Epidemiology, 29(1), pp. 158–167. Available at: https://doi.org/10.1093/ije/29.1.158.

Hastie, T. and Tibshirani, R. (1986) ‘Generalized Additive Models’, Statistical Science, 1(3), pp. 297–310. Available at: https://doi.org/10.1214/ss/1177013604.

Huang, D. et al. (2022) ‘Catalytic Priors: Using Synthetic Data to Specify Prior Distributions in Bayesian Analysis’. arXiv. Available at: http://arxiv.org/abs/2208.14123 (Accessed: 29 June 2023).

Hughes, J.H., Upton, R.N. and Foster, D.J.R. (2017) ‘Comparison of non-compartmental and mixed effect modelling methods for establishing bioequivalence for the case of two compartment kinetics and censored concentrations’, Journal of Pharmacokinetics and Pharmacodynamics, 44(3), pp. 233–244. Available at: https://doi.org/10.1007/s10928-017-9511-7.

Jaki, T., Wolfsegger, M.J. and Ploner, M. (2009) ‘Confidence intervals for ratios of AUCs in the case of serial sampling: a comparison of seven methods’, Pharmaceutical Statistics, 8(1), pp. 12–24. Available at: https://doi.org/10.1002/pst.321.

Jullion, A. et al. (2009) ‘Pharmacokinetic parameters estimation using adaptive Bayesian P-splines models’, Pharmaceutical Statistics, 8(2), pp. 98–112. Available at: https://doi.org/10.1002/pst.336.

Kay, M. (2022) ‘ggdist: Visualizations of Distributions and Uncertainty.’ Available at: https://mjskay.github.io/ggdist/.

Kneib, T., Silbersdorff, A. and Säfken, B. (2023) ‘Rage Against the Mean – A Review of Distributional Regression Approaches’, Econometrics and Statistics, 26, pp. 99–123. Available at: https://doi.org/10.1016/j.ecosta.2021.07.006.

Kümmel, A. et al. (2018) ‘Confidence and Prediction Intervals for Pharmacometric Models: Confidence & Prediction Intervals for Pharmacometrics’, CPT: Pharmacometrics & Systems Pharmacology, 7(6), pp. 360–373. Available at: https://doi.org/10.1002/psp4.12286.

Lai, T.L., Shih, M.-C. and Wong, S.P. (2006) ‘A New Approach to Modeling Covariate Effects and Individualization in Population Pharmacokinetics-Pharmacodynamics’, Journal of Pharmacokinetics and Pharmacodynamics, 33(1), pp. 49–74. Available at: https://doi.org/10.1007/s10928-005-9000-2.

Li, L. et al. (2002) ‘Estimation and Inference for a Spline-Enhanced Population Pharmacokinetic Model’, Biometrics, 58(3), pp. 601–611.

Margossian, C.C., Zhang, Y. and Gillespie, W.R. (2022) ‘Flexible and efficient Bayesian pharmacometrics modeling using Stan and Torsten, Part I’, CPT: Pharmacometrics & Systems Pharmacology, 11(9), pp. 1151–1169. Available at: https://doi.org/10.1002/psp4.12812.

Morey, R.D. et al. (2016) ‘The fallacy of placing confidence in confidence intervals’, Psychonomic Bulletin & Review, 23(1), pp. 103–123. Available at: https://doi.org/10.3758/s13423-015-0947-8.

Mould, D. and Upton, R. (2012) ‘Basic Concepts in Population Modeling, Simulation, and Model-Based Drug Development’, CPT: Pharmacometrics & Systems Pharmacology, 1(9), p. 6. Available at: https://doi.org/10.1038/psp.2012.4.

Mundo, A.I., Tipton, J.R. and Muldoon, T.J. (2022) ‘Generalized additive models to analyze nonlinear trends in biomedical longitudinal data using R: Beyond repeated measures ANOVA and linear mixed models’, Statistics in Medicine, 41(21), pp. 4266–4283. Available at: https://doi.org/10.1002/sim.9505.

Park, K. et al. (1997) ‘A Semiparametric Method for Describing Noisy Population Pharmacokinetic Data’, Journal of Pharmacokinetics and Biopharmaceutics, 25(5), pp. 615–642. Available at: https://doi.org/10.1023/A:1025769431364.

Pedersen, E.J. et al. (2019) ‘Hierarchical generalized additive models in ecology: an introduction with mgcv’, PeerJ, 7, p. e6876. Available at: https://doi.org/10.7717/peerj.6876.

Pinheiro, J.C. and Bates, D.M. (1995) ‘Approximations to the Log-Likelihood Function in the Nonlinear Mixed-Effects Model’, Journal of Computational and Graphical Statistics, 4(1), pp. 12–35. Available at: https://doi.org/10.1080/10618600.1995.10474663.

Purves, R.D. (1992) ‘Optimum numerical integration methods for estimation of area-under-the-curve (AUC) and area-under-the-moment-curve (AUMC)’, Journal of Pharmacokinetics and Biopharmaceutics, 20(3), pp. 211–226. Available at: https://doi.org/10.1007/BF01062525.

Pya, N. and Wood, S.N. (2015) ‘Shape constrained additive models’, Statistics and Computing, 25(3), pp. 543–559. Available at: https://doi.org/10.1007/s11222-013-9448-7.

R Core Team (2022) ‘R: A Language and Environment for Statistical Computing’. Vienna, Austria: R Foundation for Statistical Computing. Available at: https://www.R-project.org.

Siivola, E., Weber, S. and Vehtari, A. (2021) ‘Qualifying drug dosing regimens in pediatrics using Gaussian processes’, Statistics in Medicine, 40(10), pp. 2355–2372. Available at: https://doi.org/10.1002/sim.8907.

Simpson, G.L. (2018) ‘Modelling Palaeoecological Time Series Using Generalised Additive Models’, Frontiers in Ecology and Evolution, 6, p. 149. Available at: https://doi.org/10.3389/fevo.2018.00149.

Stan Development Team (2022) ‘Stan User’s Guide’. Available at: https://mc-stan.org/docs/stan-users-guide/.

Tian, M., Yu, J. and Kim, J. (2020) ‘Estimation of the area under a curve via several B-spline-based regression methods and applications’, Journal of Biopharmaceutical Statistics, 30(4), pp. 704–720. Available at: https://doi.org/10.1080/10543406.2020.1730871.

Van Rij, J. et al. (2019) ‘Analyzing the Time Course of Pupillometric Data’, Trends in Hearing, 23, p. 233121651983248. Available at: https://doi.org/10.1177/2331216519832483.

Vehtari, A. et al. (2021) ‘Rank-Normalization, Folding, and Localization: An Improved R^ for Assessing Convergence of MCMC’, Bayesian Analysis, 16(2). Available at: https://doi.org/10.1214/20-BA1221.

Veng-Pedersen, P. (1984) ‘Theorems and implications of a model independent elimination/distribution function decomposition of linear and some nonlinear drug dispositions. I. Derivations and theoretical analysis’, Journal of Pharmacokinetics and Biopharmaceutics, 12(6), pp. 627–648. Available at: https://doi.org/10.1007/BF01059557.

Veng-Pedersen, P. (2001) ‘Noncompartmentally-based pharmacokinetic modeling’, Mathematical Modeling of Controlled Drug Delivery, 48(2), pp. 265–300. Available at: https://doi.org/10.1016/S0169-409X(01)00119-3.

Wakefield, J. (1996) ‘The Bayesian Analysis of Population Pharmacokinetic Models’, Journal of the American Statistical Association, 91(433), pp. 62–75. Available at: https://doi.org/10.2307/2291383.

Warner, R., Ydstie, J., et al. (2020) ‘Comparative Pharmacokinetics of Meloxicam between Healthy Post-partum versus Mid-lactation Dairy Cattle’. Iowa State University. Available at: https://doi.org/10.25380/iastate.12605624.v1.

Warner, R., Ydstie, J.A., et al. (2020) ‘Comparative Pharmacokinetics of Meloxicam Between Healthy Post-partum vs. Mid-lactation Dairy Cattle’, Frontiers in Veterinary Science, 7, p. 548. Available at: https://doi.org/10.3389/fvets.2020.00548.

Wickham, H. (2016) ggplot2: Elegant Graphics for Data Analysis. Springer-Verlag New York. Available at: https://ggplot2.tidyverse.org.

Wiley, J. and Hedeker, D. (2022) ‘Package “brmsmargins”’. Available at: https://joshuawiley.com/brmsmargins/.

Willemsen, S.P. et al. (2017) ‘Flexible multivariate nonlinear models for bioequivalence problems’, Statistical Modelling, 17(6), pp. 449–467. Available at: https://doi.org/10.1177/1471082X17706018.

Wood, S.N. (2003) ‘Thin plate regression splines’, Journal of the Royal Statistical Society: Series B (Statistical Methodology), 65(1), pp. 95–114. Available at: https://doi.org/10.1111/1467-9868.00374.

Wood, S.N. (2011) ‘Fast stable restricted maximum likelihood and marginal likelihood estimation of semiparametric generalized linear models’, Journal of the Royal Statistical Society: Series B (Statistical Methodology), 73(1), pp. 3–36. Available at: https://doi.org/10.1111/j.1467-9868.2010.00749.x.

Wood, S.N. (2020) ‘Inference and computation with generalized additive models and their extensions’, TEST, 29(2), pp. 307–339. Available at: https://doi.org/10.1007/s11749-020-00711-5.

Wood, S.N. (2023) ‘Package “mgcv”’. Available at: https://cran.r-project.org/web/packages/mgcv/index.html.

Woodward, A. and Whittem, T. (2019) ‘The lower limit of quantification in pharmacokinetic analyses’, Journal of Veterinary Pharmacology and Therapeutics, 42(6), pp. 585–587. Available at: https://doi.org/10.1111/jvp.12778.

